# Unified Tumor Growth Mechanisms from Multimodel Inference and Dataset Integration

**DOI:** 10.1101/2022.07.22.500696

**Authors:** Samantha P. Beik, Leonard A. Harris, Michael A. Kochen, Julien Sage, Vito Quaranta, Carlos F. Lopez

## Abstract

Mechanistic models of biological processes can help explain observed phenomena and predict response to a perturbation. A mathematical model is typically constructed using expert knowledge and informal reasoning to generate a mechanistic explanation for a given observation. Although this approach works well for simple systems with abundant data and well-established principles, quantitative biology is often faced with a dearth of both data and knowledge about a process, thus making it challenging to identify and validate all possible mechanistic hypothesis underlying a system behavior. To overcome these limitations, we introduce a Bayesian multimodel inference (Bayes-MMI) methodology, which quantifies how mechanistic hypotheses can explain a given experimental datasets, and concurrently, how each dataset informs a given model hypothesis, thus enabling hypothesis space exploration in the context of available data. We demonstrate this approach to probe standing questions about heterogeneity, lineage plasticity, and cell-cell interactions in tumor growth mechanisms of small cell lung cancer (SCLC). We integrate three datasets that each formulated different explanations for tumor growth mechanisms in SCLC, apply Bayes-MMI and find that the data supports model predictions for tumor evolution promoted by high lineage plasticity, rather than through expanding rare stem-like populations. In addition, the models predict that in the presence of SCLC-N or SCLC-A2 cells, the transition from SCLC-A to SCLC-Y through an intermediate is decelerated. Together, these predictions provide a testable hypothesis for observed juxtaposed results in SCLC growth and a mechanistic interpretation for tumor recalcitrance.

**AUTHOR SUMMARY:** To make a mathematical model, an investigator needs to know and incorporate biological relationships present in the system of interest. However, if we don’t know the exact relationships, how can we build a model? Building a single model may include spurious relationships or exclude important ones, so model selection enables us to build multiple, incorporating various combinations of biological features and the relationships between them. Each biological feature represents a distinct hypothesis, which can be investigated via model fitting to experimental data. We aim to improve upon the information theoretic framework of model selection by incorporating Bayesian elements. We apply our approach to small cell lung cancer (SCLC), using multiple datasets, to address hypotheses about cell-cell interactions, phenotypic transitions, and tumor makeup across experimental model systems. Incorporating Bayesian inference, we can add into model selection an assessment of whether these hypotheses are likely or unlikely, or even whether the data enables assessment of a hypothesis at all. Our analysis finds that SCLC is likely highly plastic, with cells able to transition phenotypic identities easily. These predictions could help explain why SCLC is such a difficult disease to treat, and provide the basis for further experiments.

## INTRODUCTION

A mechanistic understanding of biological processes that explains causal input-output relationships and predicts population behaviors (1) remains a central challenge to all areas of quantitative biology. Mathematical models have become an established practice to specify precise relationships within a biological system, (2) and thereby hypothesize, and subsequently test, the existence of these relationships. For example, multiple mechanistic models of apoptosis execution have been formulated to explore the nature of biochemical interactions that lead to cellular commitment to death, demonstrating that careful model design and suitable data can lead to important biological insights (3–5). A more challenging situation emerges when models are formulated for biological processes that are poorly defined or understood, leading to multiple competing, and often juxtaposed mechanistic explanations for a given biological process. For example, in Small Cell Lung Cancer (SCLC), a study of circulating tumor cell-derived xenografts showed that Non-NE-related subtypes could act as a stemlike population (“source”), (6) but archetype analysis of a genetically engineered mouse model tumor showed the Non-NE subtype SCLC-Y acts as an end-state (“sink”, rather than source) (7). Therefore, continued exploration of the hypotheses generated from these works can help elucidate the differences between these and other potential explanations for tumor growth mechanisms.

This phenomenon where multiple mechanistic hypotheses are concurrently proposed but must be assessed with limited data is not restricted to quantitative biology but common to other fields with similar data availability limitations such as ecology (8) climatology (9), and evolutionary biology (10), to name a few. To address this challenge, methods such as model selection and multimodel inference have been proposed using information theoretic scoring techniques such as Akaike Information Criterion, (AIC) with limited success stemming from difficulties with model averaging (11,12) and the fact that AIC scores do not inherently describe whether a model or features within are informed by the data. More recently, the use of AI and machine learning approaches has given impetus to causal relationship inference (13) but these relationships remain difficult to elucidate, thus underscoring the need for both novel tools for hypothesis exploration, and tools that can be used with rigor in the face of missing data that may not inform all hypotheses.

The model selection process involves candidate model evaluation from a superset of plausible models, relative to a given experimental dataset. A scoring function is then applied to each model usually with the goal to identify the top-scoring model that best represents the process being investigated. A typical approach is to consider a “best” model as comprising the most relevant variables that capture important mechanistic aspects of the explored process, while excluded variables capture process features that are less relevant for the question being explored. However, variables throughout all candidate models can contribute to knowledge about the overall system (14). In the cases where data is simply less informative for a given set of hypotheses, uncertainty will remain about what constitutes a “best” model, (15), necessitating approaches such as model averaging, where parameter values can be weighted by model probability and then combined into a distribution of likely values. Unfortunately, for information theoretic applications of model averaging, this probability must be weighted and summed across all possible models, which are often not possible to enumerate exhaustively in a biological context.

To address the challenge of employing multimodel inference approaches in the context of biological processes where models cannot be exhaustively enumerated, data may not inform all model evaluation, and model averaging across all models is desired to learn about the system of interest, we introduce Bayesian multimodel inference (Bayes-MMI), a method which combines Bayesian inference with model selection and model averaging. In our approach, Bayesian model selection enables exploration of mechanistic model hypotheses by inclusion or exclusion of species and behaviors that could play a role in each biological process. Application of Bayesian principles in turn reveals the extent to which data informs a given model and its constitutive parameters.

In our system of interest, small cell lung cancer, (SCLC) we integrate the most suitable available datasets and published theories of SCLC cellular biology to enumerate mechanistic hypotheses for SCLC tumor growth. We test the resulting thousands of candidate population dynamics models via nested sampling, comparing candidate model output to tumor steady-state data, applying the principles of model selection and model averaging for a principled and comprehensive assessment of SCLC mechanistic hypotheses. We estimate the probability of each mechanistic hypothesis given the data, generating an interpretation of SCLC tumor growth: highly likely non-hierarchical phenotypic transitions indicating SCLC subtype plasticity, and less likely cell-cell interactions that affect the rate of phenotypic transitions across subtypes. We show how certain aspects of the SCLC model, such as phenotypic transitions and cell-cell interactions related to these, are well informed by the available data, but other aspects, such as tumor initiation and growth rate effects, are not informed. Our approach is generalizable to other biological systems, and as such we suggest a shift away from considering only one “best” model or the variables within and instead propose a move toward Bayesian principles in multimodel inference and a probabilistic understanding of whether each cellular behavior in a model contributes to the behavior of the system under study.

## RESULTS

### Existing datasets yield multiple hypotheses in SCLC tumor growth mechanisms

Small cell lung cancer (SCLC) has been denominated a recalcitrant tumor where relapse after treatment is commonplace and the survival prognosis is typically poor. SCLC comprises ∽15% of all lung cancer cases worldwide and results in ∽200,000 deaths annually with a 5-year survival rate of less than 10% (16). Intratumoral heterogeneity is hypothesized to be the main contributor to the natural history of this disease and its morbidity and mortality (16–18). SCLC tumors comprise a mix of functionally distinct subtypes of interacting cells, (19–21), most notably neuroendocrine (NE) and Non-NE. As shown in **Fig 1A**, SCLC populations comprise a collection of cellular subtypes within a tumor, identified by differential expression of transcriptional regulators (17).

**Fig 1.**
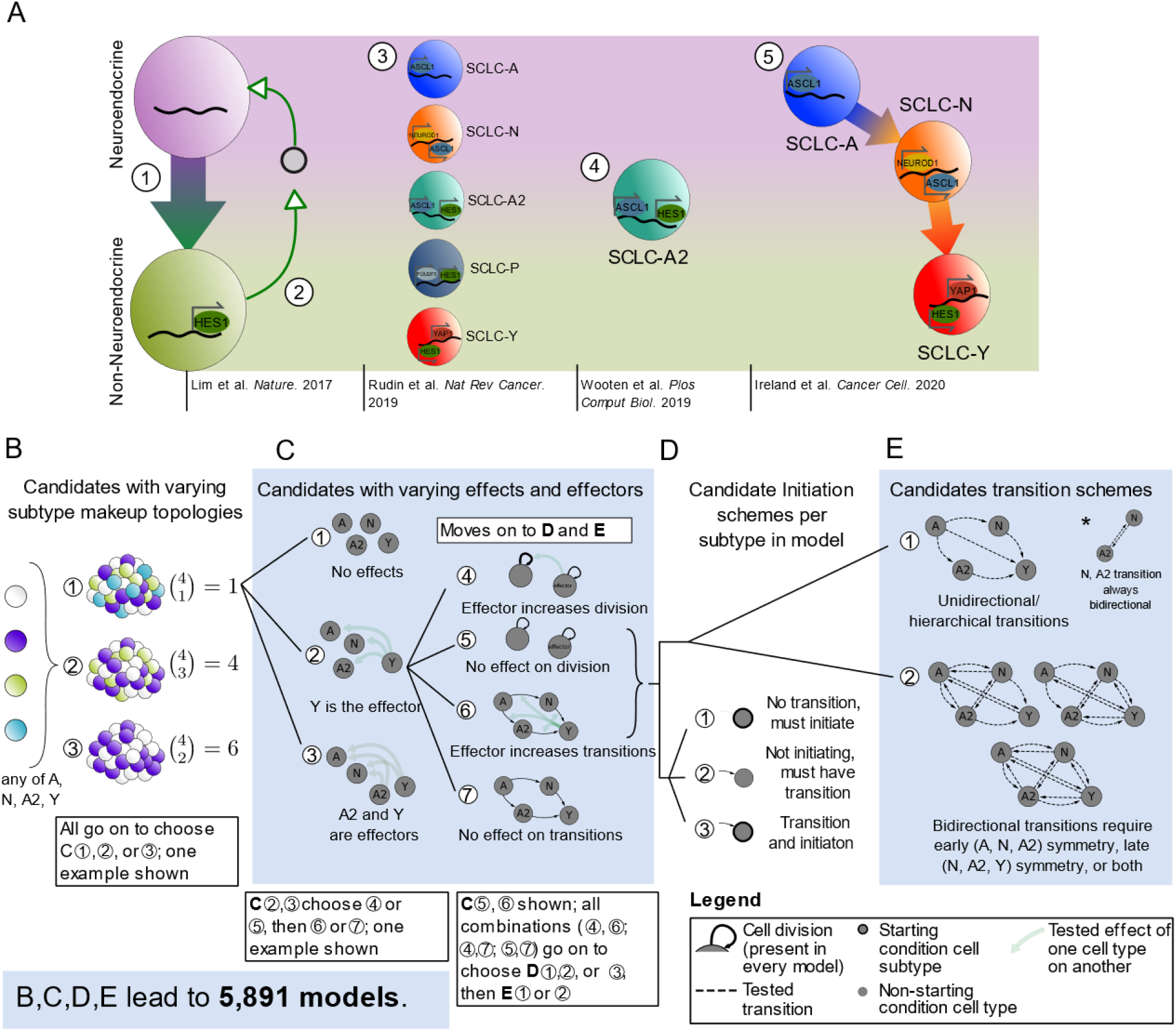
Conclusions and hypotheses from literature build mechanistic hypothesis exploration space for tumor growth and development. (A) Synthesis of what is currently known about SCLC subtypes, which have been divided into two overall phenotypes, neuroendocrine (NE) and Non-NE, and then further classified into subtypes based on transcription factor expression. (1) NE SCLC cells, which do not express Hes1, transition into Non-NE cells, which do. (2) Hes1+ cells release unidentified factors (gray circle) that support viability and growth of Hes1-cells, and the two Hes1+ and Hes1-populations grow better together rather than separately. (3) Consensus across the field led to labeling SCLC phenotypic subtypes by the dominant transcription factor expressed in that subtype. (4) Subtype with transcriptional signature intermediate between NE and Non-NE, named SCLC-A2. (5) Phenotypic transitions occur in a hierarchical manner from SCLC-A to SCLC-N to SCLC-Y cells. (B)-(E) Candidate model examples representing SCLC biological hypotheses (Table 1). (B) Model topologies constructed with 2+ subtypes, with number of combinations per number of subtypes. (C) Subtype effect schema, where there are different effectors between candidates and different affected cellular actions. (D) Potential initiation schema, where all subtypes in topology must be accessible either as initiating subtypes or via transitions. (E) Potential transition schema where, unidirectional transitions are those that follow a hierarchy, and bidirectional transitions must be symmetrical when present. HES1, Hes Family BHLH Transcription Factor 1; ASCL1, Achaete-scute homolog 1; NEUROD1, neurogenic differentiation factor 1; POU2F3, POU class 2 homeobox 3; YAP1, yes-associated protein.

The overall goal in this work is to computationally explore tumor growth mechanism hypotheses in SCLC. Tumor features that emerge as highly supported by data about the growth mechanism could be used to predict differences in growth across tumors of different genetic backgrounds, responses to *in silico* treatment, or even predict patient-specific tumor behavior after various treatments. Unfortunately, these goals are currently hypothetical, because to build one SCLC model that could be used for these purposes, one would need a unified understanding of the SCLC tumor as a system, and knowledge of SCLC currently exists as nonoverlapping conclusions and hypotheses. We summarize the current knowledge about SCLC tumor growth mechanisms, highlight potential knowledge gaps, and refer interested readers to (17) for a comprehensive review of recent SCLC literature beyond that noted here.

**Table 1.**
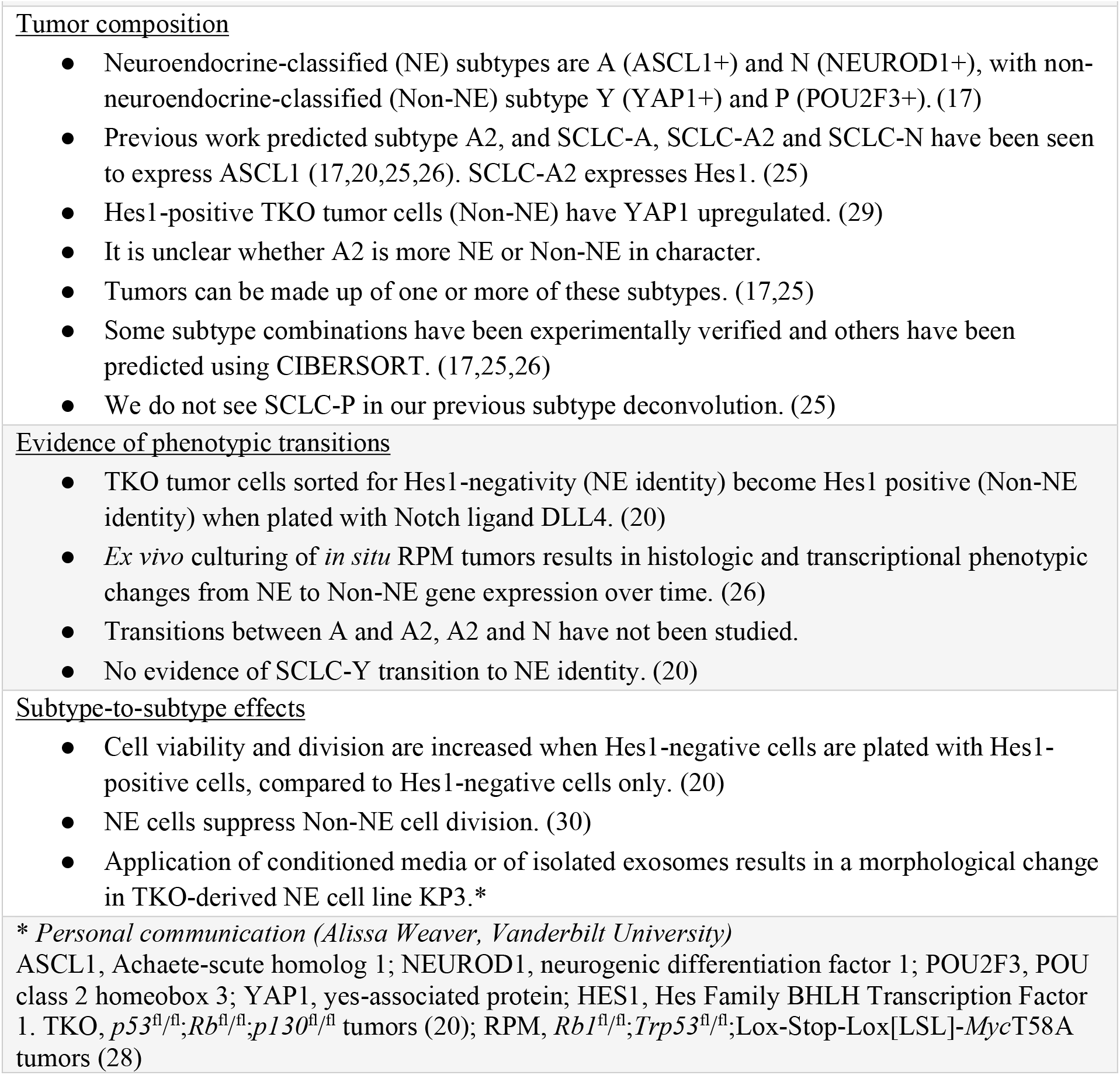
Existing data pertaining to SCLC intratumoral heterogeneity and communication.

Multiple SCLC subtypes have been identified depending on the experimental model studied, shown in **Fig 1A**, as SCLC-A, N, A2, and Y (Rudin et al., 2019). Other experiments have led to additional proposed phenotypic subtypes, including canonical subtype SCLC-P, but these were not included in our analysis (6,21–24). Our previous work aimed to identify whether all subtypes may be present in a tumor or if only a subset are present, with the result that tumors can be composed of one, multiple, or all subtypes tested (25). A comprehensive account of initiating SCLC subtype(s) (cell(s) of tumor origin) has not been made, but multiple have been hypothesized in (25–27).

Studies *in vitro* and *in vivo* have suggested that Non-NE subtype(s) support growth of NE subtypes (20) (**Fig 1A(2)**), including vasculogenic mimicking SCLC cells having such supportive effects (21). The presence of NE subtypes has a dampening effect on Non-NE growth (28). Recent work has shown that the Hes1-positive (Non-NE) cells supporting NE subtype growth (20) may have upregulated YAP1 (29) (**Fig 1A(2)**), and are likely SCLC-Y; otherwise, the referenced studies were completed before the adoption of the canonical subtypes SCLC-A, N, A2, P, and Y, and so it is unclear which of these exactly contribute such effects in each case.

NE cells may undergo a transition a toward Hes1+, (likely YAP1+) identity, (**Fig 1A(1)**) which modulates these Non-NE cells’ sensitivity to anticancer drug treatment (20). Other work found that SCLC-A subtype cells can transition to the SCLC-N subtype and from SCLC-N to SCLC-Y (26) (**Fig 1A(5)**). Without the ability to undergo a transition toward a more Non-NE phenotype, tumors were smaller and less aggressive; however, this study did not assess Non-NE or SCLC-Y sensitivity to anticancer drugs (26). These two landmark studies assessing phenotypic transitions do not assess the same phenotypic transition pathway and thus we cannot compare intermediates, although we hypothesize that the transitions begin with SCLC-A and it seems reasonable to assume the hierarchical pathway ends in SCLC-Y. While our investigations support that SCLC-Y acts as an end state for phenotypic transitions, (7) another study identified that Non-NE subtypes may have stemlike potential, (6) which contrasts with Non-NE or SCLC-Y acting as the end of the hierarchical pathway.

### Multiple mechanistic hypotheses emerge from existing data

Considering the aspects of SCLC tumor growth observed in the previous section, it is clear that no one model exists that could easily recapitulate all datasets. To address this challenge, we explored mechanistic hypotheses in the realm of tumor initiation and composition, phenotypic transitions and their hierarchy, and subtype-to-subtype effects (**Table 1**). To select which of these multiple hypotheses to include in a mechanistic model of SCLC without additional findings would introduce bias into the modeling process.

Instead, we can address these questions computationally, by including or excluding these behaviors across multiple mechanistic models and evaluating whether model behaviors recapitulate SCLC data; that is, we turn to model selection (14,31).

For our tumor growth mechanism exploration, we interpret tumor topology, initiation, potential subtype behaviors, etc., as features (model terms) in candidate models (**Table 1**). We define *model variables* as representations of species in the model (e.g., “subtype A”), and *model terms* as qualitative actions in the model (e.g., “subtype A cell division”), whose rates are denoted by *kinetic parameters* (e.g., “subtype A division rate”) (**Fig S1A-C**).

To fully account for all possible tumor subtype makeup and an exhaustive set of possible explanations, we explored models comprising between two and four subtypes per model (**Fig 1B**) and included all possible cell subtype interactions. Growth supportive effects and transition-inducing effects (**Fig 1C**) (and growth dampening effects, not shown) are included in some candidate models where, e.g., presence of an effector (supportive cell subtype) increases the rate of growth of a subtype it affects (supports). Subtype A2 has expression features of both NE and Non-NE cells (25), including expression of ASCL1 (seen in NE cells) and HES1 (seen in Non-NE cells) and we therefore assigned A2 NE features in some candidate models and Non-NE in others (**Fig 1C**).

To compare a hierarchical system, where a cancer stem cell (CSC) can (re)populate a tumor, and non-hierarchical systems in which phenotypic transitions can occur among multiple or all SCLC subtypes, we include candidate models with several different potential phenotypic transition schemes. Thus, the set of candidate models considered include models without phenotypic transitions, models with transitions that reflect hierarchical transitions observed experimentally (20,26), and models with reversible transitions, i.e., high plasticity (**Fig 1D**,**E**). Unidirectional transitions stemming from one cell subtype indicate a potential CSC, while bidirectional transitions from multiple subtypes indicate phenotypic plasticity. We additionally include tumor initiation from one cell of origin *vs*. multiple. Thus, candidate models include different numbers of initiating subtypes (**Fig 1D, Fig S2G**).

To ensure we built a comprehensive set of candidate models that enable exhaustive exploration of biologically relevant hypothesis space, we combined the potential SCLC behaviors (**Fig 1, Table 1**) with prior knowledge about mechanistic behavior of tumor populations (32–37). For example, if there is indeed plasticity in the system, it is likely to be shared among subtypes, leading to symmetrical bidirectional phenotypic transitions across the model (**Fig 1E**). We therefore expect that all plausible SCLC tumor growth mechanisms are represented in our candidate model hypothesis space to the best of our knowledge. Accounting for all these different possibilities led to a set of 5,891 unique candidate models, each representing a possible SCLC tumor growth mechanistic hypothesis.

### Bayesian exploration of candidate population dynamics models using experimental data

We use multiple datasets to identify consensus behavior of SCLC and provide a unifying model of tumor growth mechanisms broadly supported by available data (**Fig 2A**). These datasets include two genetically-engineered mouse models (GEMMs), the triple-knockout (TKO) model (*p53*^fl/fl^;*Rb*^fl/fl^;*p130*^fl/fl^ tumors (20); equivalent to the RPR2 GEMM (26,28,38)), and the RPM model (*Rb1*^fl/fl^;*Trp53*^fl/fl^;Lox-Stop-Lox[LSL]-*Myc*^T58A^ (28)), and cell lines from the Cancer Cell Line Encyclopedia (CCLE) (39) made up largely of the SCLC-A subtype determined in (25) (**Fig 2A**; **File S1**). This data provided SCLC subtype-assigned proportions of tumor samples, using the same gene signatures across samples, which had been automatically determined by CIBERSORT from samples of CCLE SCLC cell lines (39) and their consensus clustering class labels (25). We consider this preferable to our own *ad hoc* decisions of individual cell subtype identity necessary to assign the required subtype proportions of tumors had we used newer, available single-cell RNA sequencing data. The different datasets represented in **Fig 2A** demonstrate differing SCLC tumor makeup, according to the experimental model employed in the study.

**Fig 2.**
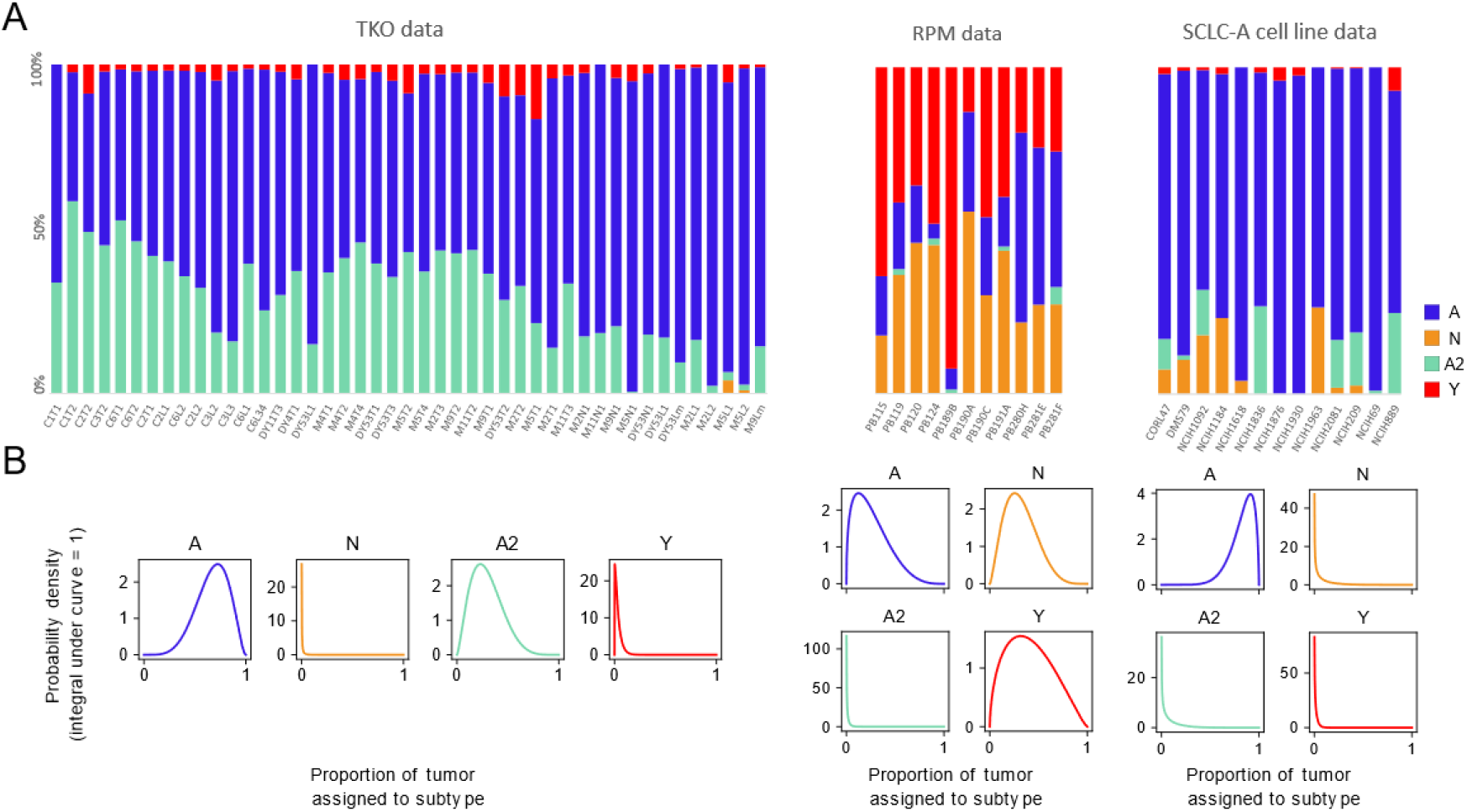
Population composition data and probabilistic representation. (A) CIBERSORT deconvolution of TKO and RPM genetically engineered mouse model (GEMM) samples (previously published) as well as SCLC-A cell line samples. CIBERSORT was performed on bulk RNA-sequencing data. (B) Probabilistic representation of tumor proportion based on mean and standard deviation of proportions across samples within an experimental model; these distributions were then used for fitting models to data. TKO, p53^fl/fl^;Rb^fl/fl^;p130^fl/fl^ tumors [20]; RPM, Rb1^fl/fl^;Trp53^fl/fl^;Lox-Stop-Lox[LSL]-Myc^T58A^ tumors [28]; SCLC-A cell lines, a subset of SCLC cell lines from the CCLE [39] that we previously assigned as representative of tumors made up largely of the SCLC-A subtype [25].

To explore the roles of phenotypic heterogeneity and cellular behaviors (cell-cell interactions, phenotypic transitions) on SCLC tumor growth dynamics, we used population dynamics modeling, building on our previous work (40). Population dynamics models employ a mathematical description of the dynamics within and between heterogeneous subpopulations in an overall population (41,42). With such models, researchers can mathematically simulate population growth over time and investigate growth dynamics inherent in the simulations (**Box 1**).

With 5,891 candidate models, (**Fig 1B-E**) we aimed to determine which one(s) could best represent the SCLC system – in an ideal world, we would determine which model represents reality. However, because we will never know the “true” model, we must evaluate each model probabilistically – which model is most likely to explain the data? Before comparing our 5,891 models to data to determine which is best, we consider each to have the same probability of explaining the data. That is, we assign all models an equal probability, as this way, we will not bias the exploration of the data toward any particular model or group of models. This is the reason we built a set of models that enables full exploration of the hypothesis space, (detailed above) so that with the addition of data, we can ensure we will study all possibilities and find the best model(s) within that hypothesis space. We compare each candidate model to the data discussed above, and with model optimization, we assess how well its simulations match what we see in that data. After this, each model’s probability will be updated, with a model that explains the data better achieving a higher probability than one that does not explain the data as well.

The principles of model selection enable us to assess which candidates are best supported by experimental data, while prioritizing model simplicity (14,43). Information theoretic approaches for model selection using the Akaike Information Criterion (AIC) have been used in prior work but do not yield a much-needed statistical understanding of the data. For a didactic demonstration of this, we refer the reader to **Note S1**. We therefore employed the marginal likelihood as a more principled means for model ranking and model averaging.

#### Box 1.

**Population dynamics modeling and inter-subtype effects**

A population dynamics model represents behaviors of species over time, tracking the size of a cell population, rather than tracking individual cells. However, such a model can also include signaling and dependence between species (Harris et al., 2019). Cell population changes are represented as reactions, where subpopulation abundances increase or decreases due to varying events (division, death, phenotypic transitions), and the rates of increase or decrease can be affected by the presence of other subpopulations. Here, cell types *x* and *w* are able to undergo division, death and phenotypic transition at rates *k*_*div*_, *k*_*die*_, and *k*_*x*-*w*_, respectively (A).

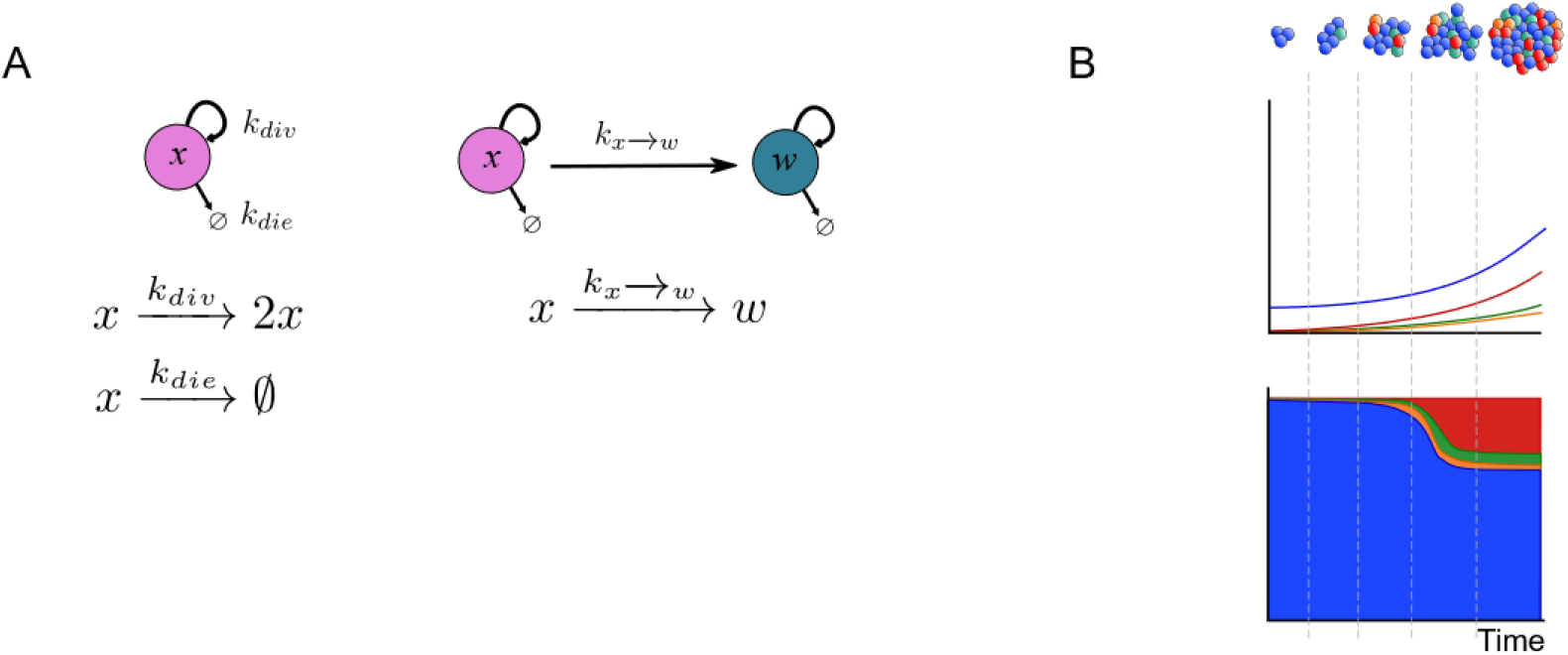

In our case, where species are cells of an SCLC tumor population, each is assigned a subtype identity (A N, A2, or Y). The model simulates tumor growth over time, (B, top) calculating the increase in subpopulation amount (B, middle), from which tumor subtype proportion may be calculated (B, bottom) and compared at steady state to tumor proportion data. We define steady state as the composition of a tumor based on the relative abundance of each cell subtype in the tumor, without external perturbations. Though tumor growth may continue exponentially in a steady state, the proportion of subpopulations within remains constant (Harris et al., 2019).

Non-spatial cell-cell interactions may also be modeled, by changing the rates of reactions for one subpopulation according to the amount of another subpopulation present. Here, using the hypothesized biology where subtype wproduces a secreted signal *f* that affects subtype *x*, we calculate the change in reaction rates. In (C), *w* secretes the unknown signaling factor *f* at a rate of *k*_*f*_, while *f* is degraded at rate *k*_-*f*_ Factor *f* may affect subtype *x* at a rate of *k*_*e*_, where *x* becomes *x*^*^ (“*x* under effect of *f*”). Subtype *x* can then revert back to its unaffected form at a rate of *k*_-*e*_, indicating the rate at which the signaling is completed.

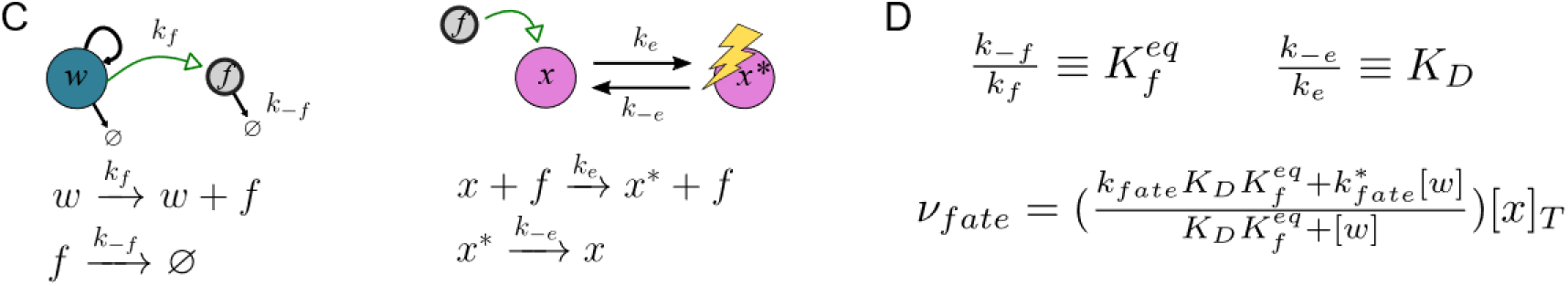

With *k*_*f*_ production rate constant and *k*_-*f*_ the degradation rate constant,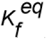 is the equilibrium constant for the amount of factor *f* in the system (D). Similarly, *K*_*D*_ is the equilibrium constant related to the on-effect rate constant *k*_*e*_ and the off-effect rate constant *k*_-*e*_. Given these, the rate of a cell fate (division, death, or phenotypic transition) for *x* (*v*_*fate*_) can be calculated as a function of the population size of the effector cell *w*(D) (Harris et al., 2019) (see Note S1 for calculations).

By assigning the value of 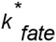 as more or less than *k*_*fate*_ for a particular cell fate, the presence of the effector cell subpopulation can increase or decrease, respectively, the rate of the cell fate for subpopulation *x*. In our population dynamics model, typically effector cells increase division and transition rates and decrease death rates, as shown in (E).

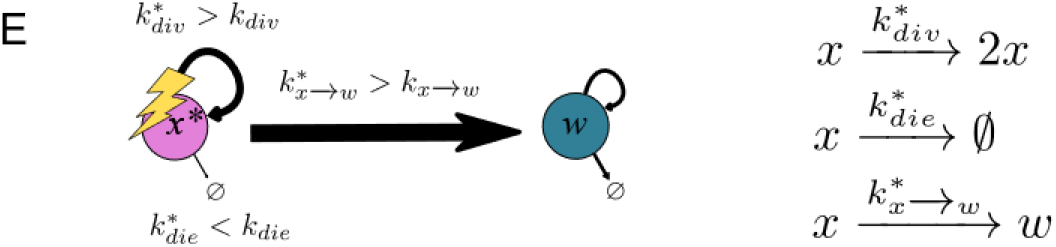

Prior work has estimated the marginal likelihood for kinetic model fitting using thermodynamic integration (44). In this work we instead use nested sampling, (45), which is computationally more efficient and has fewer limitations with regard to the shape of the probability space traversed during evidence calculation (46–48). The nested sampling method was run once for each of the 5,891 candidate models on each of the three experimental datasets, amounting to 17,484 potential interpretations of tumor growth mechanisms. The average fitting time for each model was ∽19 wall-clock hours, thus necessitating high-performance computing for a complete parameter space exploration of the candidate models.

Each model is thus optimized to our datasets via nested sampling, which explores the full volume of the likely parameter space. Each point in parameter space represents a set of possible parameter values (**Fig S3**). At each of these points, nested sampling assigns a likelihood value for how well that set of parameter values fits the data. On completion of the algorithm, the output includes the highest-likelihood parameter values. Since each tested point in parameter space is a set of parameter values, the highest-likelihood values for a model are returned by the algorithm as a list of parameter sets (**Fig S4**). Returning a list of parameter sets rather than one top-scoring set already incorporates Bayesian methodology into the process – each individual parameter has multiple best-fitting values, which can be interpreted as a distribution of parameter values (44) – but with nested sampling we add yet more Bayesian methodology. Having assigned a likelihood to every point in parameter space, nested sampling uses these to calculate one overall likelihood per model, the marginal likelihood, which takes into account parameter fit as well as model simplicity (number of parameters). For more detail on how the marginal likelihood is calculated to incorporate both model fit and size, see **Note S1** and **Methods**. Finally, with marginal likelihood values for each model in the candidate set, and the candidate model set representing the full hypothesis space with all potential SCLC population dynamics models, we can calculate a probability. Summing the marginal likelihood values, and dividing each individual marginal likelihood by these, results in a model posterior probability, representing a change in probability from pre-model fitting (all models with equal prior probability) to post-model fitting (see **Note S1**). We are then able to compare model probabilities and additionally perform model averaging to evaluate kinetic parameter value distributions and probabilities of model variables and terms.

### A small subset of candidate tumor growth models is supported by experimental data

In a Bayesian model selection approach, a more likely model comprises a higher proportion of the probability of the candidate model space (**Fig 3A-C**). After nested sampling, our results indicate the highest-scoring model for each dataset is ∽10^19^ times more likely than the lowest-scoring model, and ∽10^3^ times more likely than the median scoring model. For reference, the smallest comparison between models that is considered significant is 10^1/2^ (49) (**Note S1**). Therefore, our results indicate that the data used for model fitting has informed our knowledge about the system, because before nested sampling, all models are equally likely.

**Fig 3.**
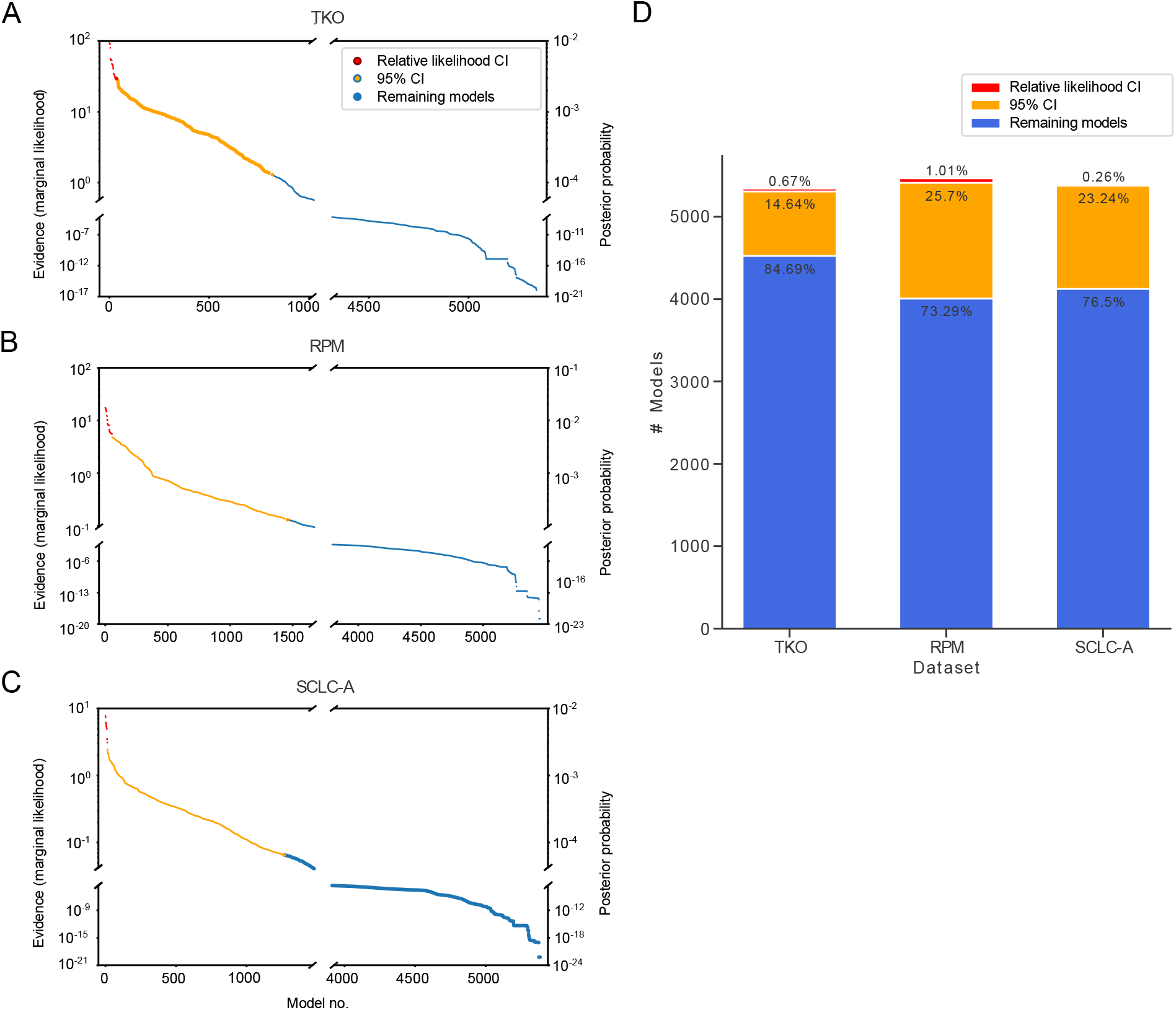
Fitting to data and assigning Bayesian evidence separates candidate models into more and less likely. (A)-(C). Evidence values (left y-axis) and posterior probability values (right y-axis) from nested sampling, one point per model, ordered from model with greatest evidence to model with least evidence. Models whose evidence value are within 101/2 of the greatest evidence value, the “relative likelihood confidence interval,” are colored in red. Nested sampling and evidence calculation is performed per dataset. (A) TKO dataset. (B) RPM dataset. (C) SCLC-A cell line dataset. (D) Numbers and percentages of models in the relative likelihood confidence interval, 95% confidence interval, and remaining non-confidence interval models. TKO, *p53*^fl/fl^;*Rb*^fl/fl^;*p130*^fl/fl^ tumors [20]; RPM, *Rb1*^fl/fl^;*Trp53*^fl/fl^;Lox-Stop-Lox[LSL]-*Myc*^T58A^ tumors [28]; SCLC-A cell lines, a subset of SCLC cell lines from the CCLE [39] that we previously assigned as representative of tumors made up largely of the SCLC-A subtype [25].

Performing nested sampling on all candidate models did not yield a unique best-fitting model for any dataset (**Fig 3A-C**). We therefore leveraged a multi-model inference approach and calculated a confidence interval (CI) representing a set of best-fitting models per dataset. While a 95% CI is a traditional approach, (**Fig 3A-C**, orange) we also calculate a “relative likelihood confidence interval,” as discussed in (14) (see **Methods**). For this relative likelihood CI, we calculate the Bayes Factor (BF) between the highest-scoring model and every other model, using the least strict cutoff of BF > 10^1/2^ (as above, and in **Note S1**). Even with this permissive cutoff, the relative likelihood CI includes only tens of models per dataset, a large decrease from the initial number of candidates (∽1% or less, **Fig 3D**).

In summary, we can determine a subset of candidate models that adequately represent the data, conditional on the fitted parameter sets resulting from the model optimization in nested sampling. Investigating these parameter sets can provide more insight into the similarities and differences between candidate models and their fits within and between datasets. Moving beyond the parameter values assigned to each model term, we wanted to investigate how the data available can inform model terms. If data does not inform model terms and variables and the corresponding fitted parameter rates, it indicates that the mechanistic conclusions we desire to draw from this data using mathematical modeling may require additional or different data.

### High-likelihood model topologies are nonoverlapping between datasets

Given our observation that no one candidate model stands out among other models to explain the experimental data, we employed the multi-model inference technique of Bayesian model averaging (BMA). Briefly, the reasoning behind BMA is that a combination of candidate models will perform better in explaining the data than a single model (15). In BMA, each model is weighted by its posterior model evidence (31) and the model terms within each model receive an averaged likelihood (50) (**Note S1-S3**).

However, before we could investigate model-averaged probabilities, we found that the fitted parameter distribution outcome was dependent on choice of initiating subtype (**Fig S5**). While we aimed to evaluate initiating subtype possibilities with our Bayes-MMI approach, we cannot do this in a unifying fashion as this choice affects fitted parameters and thus model behavior. Therefore, we turned to the literature to impose stricter constraints about initial subtype conditions, to investigate model variable and term probabilities when kinetic parameter posterior distributions were constrained. As mentioned previously, reports link NE SCLC subtypes and long-term tumor propagation (20,30) and, in particular, cells of subtype A (26). We thus used only candidate models with an initiating subtype of A, with or without other initiating subtypes. Since we required that subtype A be an initiating subtype, model structures that do not include subtype A received zero posterior probability (models 3 and 8–10 in **Fig 4A**; model topology probabilities without filtering by initiating subtype are shown in **Fig S6A**).

**Fig 4.**
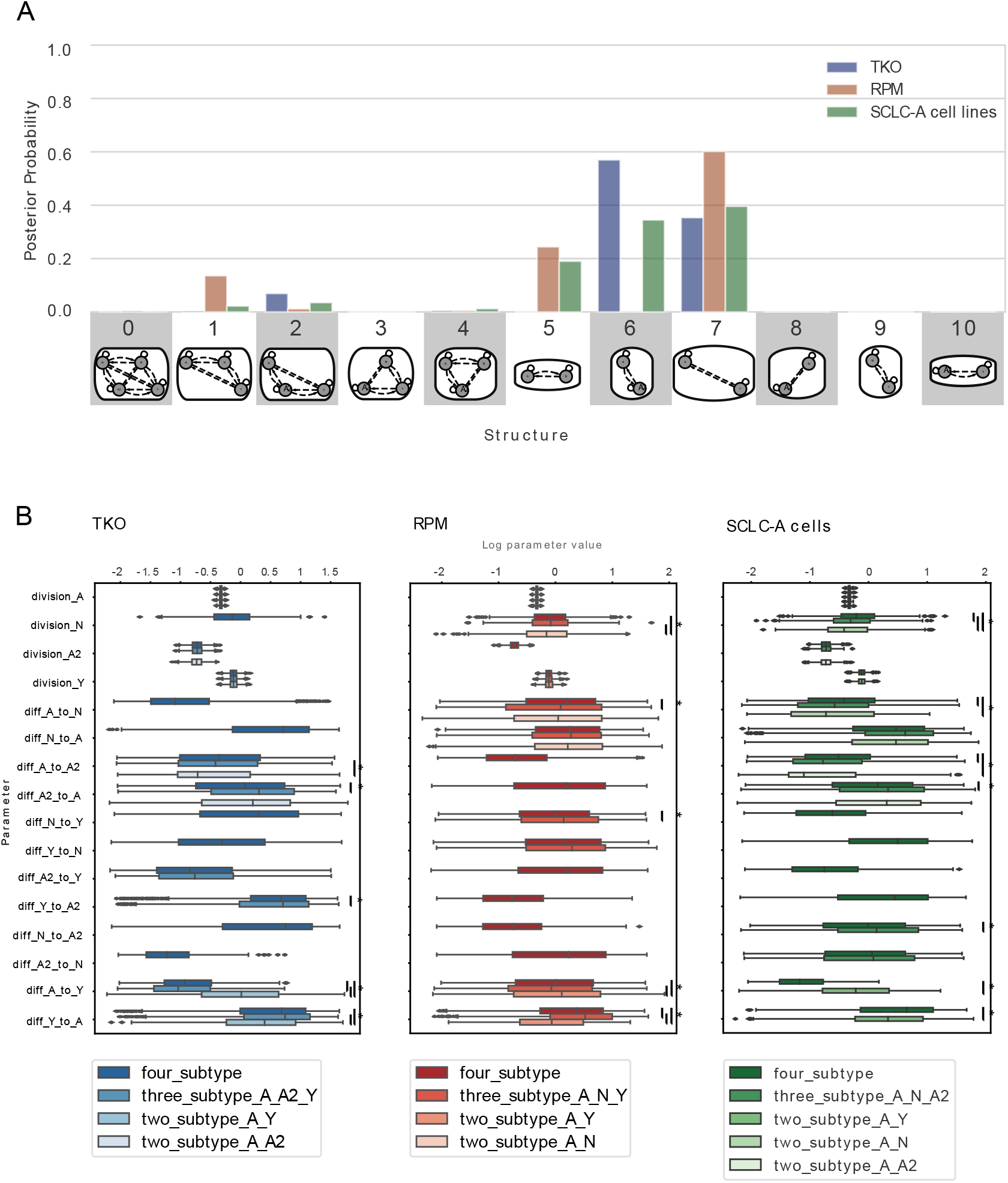
Likely model topologies vary across datasets; transition rates vary according to subtype presence in similar ways. (A) Hypothesis assessment of model topologies, per dataset. Probability indicates the result of Bayes theorem using equivalent prior probabilities per topology (e.g., 9% probability that one of the topologies in the x-axis best represents a dataset) and evidence values (marginal likelihoods) summed per topology. Model topologies represented by images and corresponding numbers along the x-axis. Posterior probability based on marginal likelihoods of all candidate models that include A as an initiating subtype. (B) Division and phenotypic transition parameters for TKO, RPM, and SCLC-A cell line datasets, comparing between higher-probability topologies (A) and four-subtype topology per dataset. TKO, *p53*^fl/fl^;*Rb*^fl/fl^;*p130*^fl/fl^ tumors [20]; RPM, *Rb1*^fl/fl^;*Trp53*^fl/fl^;Lox-Stop-Lox[LSL]-*Myc*^T58A^ tumors [28]; SCLC-A cell lines, a subset of SCLC cell lines from the CCLE [39] that we previously assigned as representative of tumors made up largely of the SCLC-A subtype [25]. (*) indicates significance between samples from BMA parameter distributions at family-wise error rate (FWER) = 0.01, averaged over ten sampling iterations using one-way ANOVA plus Tukey HSD.

We perform BMA across all models for each dataset. As shown in **Fig 4A**, all datasets (TKO, RPM, and SCLC-A cell lines) support both two- and three-subtype topologies. Higher probabilities for two-subtype topologies are expected given that nested sampling prioritizes model simplicity and goodness of fit (45). Statistically, this result suggests that a two-subtype model could be used to interpret the data reasonably well, but it also shows that topologies comprising three subtypes cannot be excluded. Approximately 10% of the probability for the GEMM datasets (TKO and RPM) fall in the three-subtype topology that encompasses the high-probability two-subtype topologies (model 1 for RPM and model 2 for TKO in **Fig 4A**). For the models fit to SCLC-A cell line data, most of the probability occurs in the topologies with higher probabilities for the GEMM data. This is reasonable, given that SCLC-A cell line data appears as an intermediate between the GEMMs (**Fig 2A**). However, the SCLC-A cell line data also has probability that falls in the A, N, and A2 topology (model 4 in **Fig 4A**) – this is the only topology at all likely to represent the SCLC-A cell line data but not at all likely to represent the other two datasets.

We interpret the spread of probabilities across multiple topologies, and that most topologies either are probable as representing either TKO or RPM data but not both, to mean that data coverage from these datasets is not sufficient to support one unifying topology. Therefore, each dataset supports a different representation of SCLC tumor growth given its particular (epi)genetic background and environment. This does not mean that a unifying topology or unifying model of SCLC growth cannot exist, but that the biases underlying the experimental data result in different explanations for tumor growth mechanisms.

### All datasets support alteration of phenotypic transition rates in the presence of N or A2 subtypes

After establishing that multiple model topologies can explain tumor growth mechanisms, we explored whether the rates of distinct cellular subtype behaviors were characteristic for given model topologies within each dataset. We therefore compared kinetic parameters across models to learn about dynamic variation between model topologies. We again used BMA to attain this goal, applying the approach to fitted kinetic parameter distributions from nested sampling. In this setting, parameter values from more likely models are assigned higher weights and corresponding parameter distributions are weighted accordingly.

The highest likelihood model topologies (**Fig 4A**, blue) for the TKO GEMM data, along with the four-subtype topology, are compared in **Fig 4B** (left). Three model terms have significantly different parameter rates across model topologies, all of which are discussed in **Note S4**. Here we highlight differences in the A-to-Y and A-to-A2 transitions across model topologies in the TKO dataset: the A-to-Y transition has a slower rate if A2 is present in the population, and only Y affects the A-to-A2 transition, increasing its rate. The mechanistic implication of these observations are as such: A2 may represent an intermediate subpopulation in the tumor that is longer-lived, and will only slowly transition to Y. In the topology with A, A2, and Y (**Fig 4A**, structure 2; **Fig 4B** left, third-darkest blue), the A-to-A2 transition takes up more of the flux in the network. Additionally, the N-to-Y transition is faster relative to the A2-to-Y transition (**Fig S6B**), suggesting that N is a shorter-lived intermediate in the A-to-N-to-Y transition. This result aligns with previous experiments (Ireland et al., 2020) where N was identified as a short-lived state in the A-to-N-to-Y transition. We therefore predict that A2 and N are involved in regulating the relative abundance of, and flux between, A and Y in the tumor.

We also compared the highest likelihood model topologies (**Fig 4A**, red) for RPM-fitted models, as well as the four-subtype topology (**Fig 4B**, middle). Five parameter rates are significantly different across model topologies, and we highlight again the A-to-Y transition and here the A-to-N transition (see **Note S4** for discussion of the remaining significantly different parameter rates). The same A-to-Y transition affected in the TKO-fitted models is affected in the RPM model in the same way (reduced rate via an intermediate, in this case N), and the A-to-N transition is affected by Y similarly to the A-to-A2 transition affected by Y in the TKO data. These similar effects occur despite the experimental data used for BMA being different. We thus predict that N and A2 are modulating the transition between, and relative abundance of, A and Y. Unlike in the TKO data, when A2 is present the flux through the system spends more time in the N subtype, with more frequent transitions to N and less frequent transitions to Y. We predict that while N may be a shorter-lived intermediate than A2, A2 regulates the flux from A-to-N-to-Y.

Next, we compared the highest likelihood model topologies (**Fig 4A**, green) for the SCLC-A cell line data and the four-subtype topology (**Fig 4B**, right). Six parameter rates are significantly different across model topologies, five of which recapitulate rate alterations based on the presence or absence of different subtypes in TKO or RPM datasets, including the rate alterations discussed above (see **Note S4** for more detail).

In summary, BMA enabled us to determine that the A-to-Y transition is regulated in a similar manner for the RPM, TKO, and SCLC-A datasets and that the A-to-N and A-to-A2 transitions are regulated similarly in each dataset. Using the higher likelihood model topologies and model-averaged parameter sets, we can infer features of the SCLC tumor generally, despite disparate datasets. Finding the same or similar effects on kinetic parameter rates across independent datasets lends more weight to these predictions about how the N and A2 subtypes may regulate the system flux from A to Y through intermediates and is an advantage of our methodology using Bayes-MMI to work toward a unifying model of SCLC tumor growth based on multiple datasets.

### Model analysis supports a non-hierarchical differentiation scheme among SCLC subtypes

We have considered candidate models (**Fig 3**), model topologies (**Fig 4A**), and kinetic parameters (**Fig 4B**) to explore tumor growth mechanisms in SCLC. There is compelling experimental evidence for multi-subtype tumor composition, which implies multiple potential growth mechanisms (6,20,26,28,51). We therefore focused on model topologies 1, 2, and 4, which are three-subtype topologies with detectable probability (> 1%) (**Fig 4A**), along with the four-subtype model. Using these, we integrate candidate models, topologies, and kinetic parameters, investigating phenotypic transitions between subtypes, whether the presence of certain subtypes affects the behaviors of other subtypes, and if so, which subtypes bring about the effects (**Table 2**). We conclude by proposing a unifying four-subtype model of tumor growth in SCLC, aiming to represent with one model the varying growth mechanisms accessible across datasets.

**Table 2.**
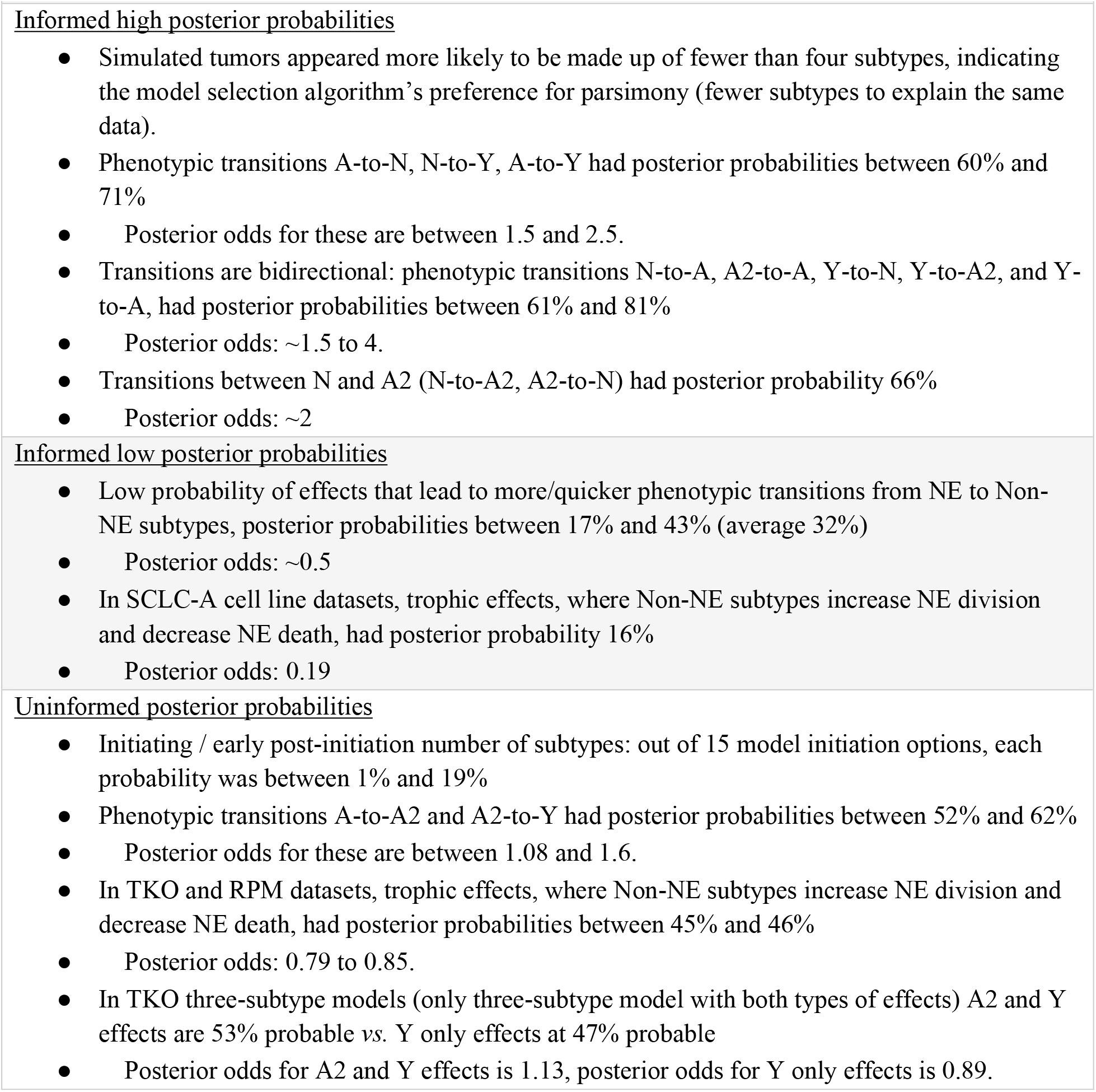
Probabilities after hypothesis exploration using Bayesian multimodel inference.

We investigate the posterior probabilities, and therefore posterior odds, of each model term (see **Methods**). Despite different posterior probability values (**Fig 5A**), the probabilities of model terms across datasets were similar in their trends: across all three-subtype topologies, phenotypic transition probabilities were all more than ½ (**Fig 5A**, red squares). While some probability values were poorly informed (light red), (probability between ½ and ⅔), more were informed by the data (deep red) (⅔ or more). Conversely, probabilities of Non-NE effects on the growth or transitions were all less than ½ (**Fig 5A**, blue squares). Some probability values were poorly informed, (light blue) (between ⅓ and ½) and others were informed (deep blue) (⅓ or less) with the addition of data.

**Fig 5.**
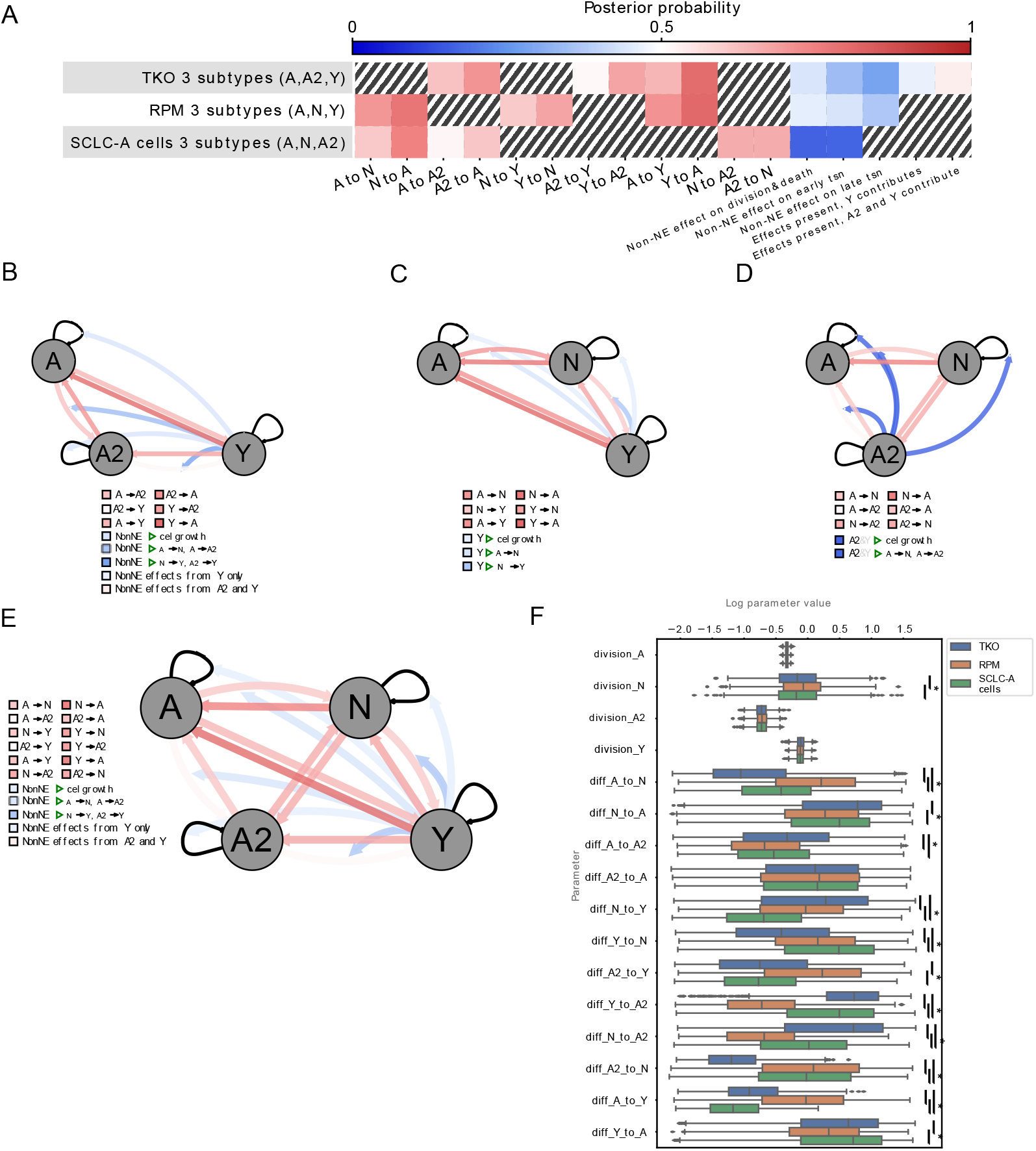
Across datasets, multimodel inference indicates likely bidirectional phentoypic transitions, suggesting high SCLC phenotypic plasticity. (A) Heatmap for high probability three-subtype topologies for each dataset (rows), all models initiated by A +/-other subtypes. Color represents the probability of each cellular behavior (column). Since prior probability starts at 0.5 (white), deeper colors indicate a larger deviation from the prior, with red vs blue indicating more likely or less likely, respectively. (B)-(D). Model schematics with each cellular behavior represented by edges coming from or moving toward each cell subtype, (gray circles) growth rates, (self-arrows) or transitions (arrows between gray circles). Edge colors correspond to colors for that behavior in the heatmap in (A). Top-scoring three-state topology for TKO dataset (B), RPM dataset (C), and SCLC-A cell line dataset (D). (E) Schematic of consolidated model behaviors, drawn from each dataset’s high-probability three-subtype topology results ((B)-(D)). When multiple dataset results included different posterior probabilities for a model feature, the one closest to 0.5 was chosen (most conservative). Edge colors correspond to posterior probabilities, with intensity of colors representing information gained from data, as in (A)-(E). (F) Parameter fitting results (part of the nested sampling algorithm) for four-subtype topology models initiated by A +/- other subtypes, across datasets. tsn, transition (e.g., subtype transition). TKO, *p53*^fl/fl^;*Rb*^fl/fl^;*p130*^fl/fl^ tumors (20); RPM, *Rb1*^fl/fl^;*Trp53*^fl/fl^;Lox-Stop-Lox[LSL]-*Myc*^T58A^ tumors (28); SCLC-A cell lines, a subset of SCLC cell lines from the CCLE (39) that we previously assigned as representative of tumors made up largely of the SCLC-A subtype (25). (*) indicates significance between samples from BMA parameter distributions at family-wise error rate (FWER) = 0.01, averaged over ten sampling iterations using one-way ANOVA plus Tukey HSD.

Overall, the data suggests that Non-NE effects on transition rates of N-to-Y, or A2-to-Y, are unlikely, (**Fig 5B-D**, deep blue) regardless of whether “Non-NE” defines only the Y subtype, or both A2 and Y are Non-NE (**Fig 1C**). Inter-subtype effects on SCLC phenotypic transition rates have not previously been studied and our analysis predicts that at least effects on “late transition” (**Fig 5A**), those interactions affecting N-to-Y or A2-to-Y, are unlikely to exist. By contrast, transitions involving A-to-N, N-to-A, A2-to-A, N-to-Y, Y-to-N, A2-to-Y, Y-to-A2, N-to-A2, A2-to-N, A-to-Y, and Y-to-A had posterior probabilities informed by the data (**Fig 5B-D**, deeper red). We interpret these results as transitions being likely, i.e., our degree of belief in these transitions has increased. Investigating initiating events via one or multiple cells of origin across the candidate models, we find that from equal prior probabilities of 6.67% per initiating subtype(s) (**Fig S2G**) the posterior probabilities are not significantly altered, being between 1% and 15% (not shown). Thus, initiating subtype events were poorly informed by the data. Additionally, analyzing specific model terms, inter-subtype effects on NE subtype growth, inter-subtype effects on transition rates between A and N, or A and A2, and the A-to-A2 transition, were also poorly informed by the data (**Fig 5B-D**, light blue, light red).

Finally, to consolidate phenotypic transitions and cell-cell interactions into a unifying mechanism for SCLC tumor growth, we integrated model probabilities from each of the three-subtype topologies for each dataset into one model (**Fig 5E**). Briefly, phenotypic transition probabilities were chosen from the models least informed by the data in an attempt to make conservative predictions (see **Methods**). Model-averaged parameter rates were visually compared (**Fig 5F**) to ensure that they were within reasonable bounds and that transition rates relate to each other between datasets similarly to our analyses using high-probability topologies (**Fig 4B**). Values from the consolidated probabilities (**Fig 5E**) are those reported in **Table 2**, along with posterior odds to compare one hypothesis to its opposite.

Taken together, these results provide insight not only into what model terms and variables the data is able to inform, but SCLC tumor behavior as well. Knowledge of trophic effects provided by Non-NE cells to the benefit of NE cells was not provided by this particular data; therefore, we cannot use it to understand this behavior. However, we were able to gain knowledge about the likelihood of phenotypic transitions, in fact indicating that nearly all options for phenotypic transitions are likely to exist. We interpret this as high SCLC plasticity, supporting a non-hierarchical differentiation scheme where tumor population equilibrium is achieved through any phenotypic transitions (**Fig 5E**). It is also clear that consolidating the results across different tumor types is an important step in order to achieve a broader view of the SCLC tumor as a system rather than as one particular experimental model.

## DISCUSSION

The experimental data used for this analysis favors two-subtype topologies as higher-probability candidates. This is not surprising, because nested sampling prioritizes simpler models. Despite this finding, mounting experimental evidence supports multi-subtype tumor composition and these data have been interpreted in the context of multiple existing phenotypes (6,20,26,28,51). In fact, previous work from our labs suggests that the tumor genetic background dictates possible phenotypic subtypes within a tumor population, and that phenotypic transitions likely mediate the ability of tumor cells to achieve these phenotypes (40,52,53). We hypothesize that with additional timecourse and perturbation data, the topology likelihood for tumor growth mechanism could be shifted toward three- or four-subtype models. This is consistent with studies of tumor dynamics from cancer broadly, both with regard to phenotypic transitions toward or away from a rarer subpopulation (34,54–56) and changes in the proportion of phenotypic subpopulations after a perturbation. Often this perturbation is the application of drug treatment (57–62), but may also be changes in microenvironment or related factors (26,63).

The results presented here also provide strong evidence for phenotypic plasticity in SCLC tumors, based on the higher likelihood for most phenotypic transitions tested, regardless of differentiation hierarchy. With a more plastic and less stem-cell based phenotypic equilibrium, instead of rare remaining stem-like populations leading to regenerate a tumor after treatment, we hypothesize that any SCLC subtype that remains post-treatment can lead to tumor regeneration and subsequent treatment resistance, patient morbidity and mortality. While hierarchical phenotypic heterogeneity *vs*. phenotypic plasticity has not been experimentally studied in SCLC, we predict that plasticity is highly likely, and extremely important in the growth and evolution of the SCLC tumor. It is of particular interest to compare phenotypic plasticity and the prevalence of non-hierarchical transitions in treated *vs*. untreated tumor samples, as treatment is likely to alter the mechanisms by which tumor population equilibrium is maintained. Time-course experiments with surface marker labeling or live-reporter imaging can resolve and provide confirmation for bidirectional phenotypic transitions, which are crucial to understand in order to battle SCLC treatment resistance.

The invasive or metastatic potential of the SCLC tumor is known to be increased by Non-NE subtypes (21,64). It is unclear whether the conclusions and predictions presented here apply to SCLC in the invasive or metastatic setting, but future work will include model additions to place the tumor growth in a physiologic context that includes both the tumor *in situ* and during invasion.

We believe a shift from information theoretic multimodel inference toward a Bayesian approach, enabling investigation of optimal model(s) with identifiable parameters for a particular dataset, will benefit modeling in systems biology. The methodology employed herein incorporates model selection and model averaging into a multimodel inference framework, followed by Bayesian analysis to identify not only whether a hypothesis investigated via mechanistic modeling is or is not likely, but *how* likely (and thus how informed by the data) that hypothesis is. Understanding which hypotheses are informed by the data is especially important given variability between data in investigations of the same systems, such as a particular tumor type. It is difficult to attain a consensus model since investigators use varying experimental models within the same physiologic or disease process and thus may draw nonoverlapping conclusions, building parts of a picture but not a whole. Striving for the whole picture, via principled statistical analysis, to be followed by experiments based on informed model predictions, will advance cancer research and lead to better treatments.

## METHODS

### CIBERSORT deconvolution of RNA sequencing data

Data from two GEMM models provide multiple replicates of tumors from two genetic backgrounds: one from *p53*^fl/fl^;*Rb*^fl/fl^;*p130*^fl/fl^ (triple-knockout, or TKO) GEMM tumors (20), and another from *Rb1*^fl/fl^;*Trp53*^fl/fl^;Lox-Stop-Lox[LSL]-*Myc*^T58A^ (RPM) GEMM tumors (28). We also used publicly available SCLC cell line data. Having been originally derived from human tumors, each cell line has a different genetic background, and therefore we have only one (genetically identical) replicate per cell line sequencing event. To approximate genetic similarity between cell lines, and thus approximate multiple replicates, we expect that cell lines exhibiting similar steady state composition will be more genetically similar than those whose steady state compositions differ. Previously, in (25), we both clustered publicly available SCLC cell line data into clusters that align with the different SCLC subtypes and used CIBERSORT to deconvolute the proportions of cell line data and tumor samples into SCLC subtypes from their RNA sequencing signatures. Results used for this publication can be found as **File S1**.

### Population dynamics modeling in PySB

A population dynamics model represents the abundance of species over time, whether increase or decrease due to birth/growth or death. We use ordinary differential equation (ODE) models coded via PySB to generate population dynamics models (65). PySB is a rule-based modeling language, where one will encode

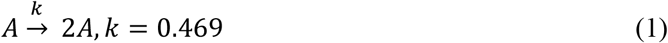

to indicate that A doubles at a rate of 0.469 doublings per day.

Inter-subtype effects are represented by the increase or decrease of the rate of affected reaction. For example, the above division rule has a baseline rate of 0.469 doublings per day, but in the presence of an effector subtype the division rule will have a rate of 0.469*1.05 = 0.493 doublings per day. In this case the effector subtype has increased the division rate by 5%. Thus the rule-based representation is

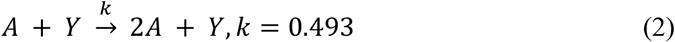

to indicate that A doubles at a rate of 0.493 doublings per day in the presence of Y.

To simulate the passage of time, the speed at which the division/death/transition reaction occurs–its rate, *k*, of cells per unit time–must be assigned as in the equations above. While a literature search reveals approximate rates of division and death among different SCLC subtypes (**Table S5**), each of these are in a different context than the system we model here – for example, division rates for the A subtype are measured *in vitro* in the presence of only that one subtype, whereas our population dynamics model is meant to simulate this subtype in the presence of others as well as *in vivo* in a mouse tumor. Therefore, we use the rates in the literature as our prior expectations for division and death, that is, we use these values as approximate starting values for these parameters during the estimation process. Other rates, such as those indicating the speed of transitions between subtypes, or any rates including the effects of Non-NE subtypes, have not previously been noted in the literature and we used much wider ranges for each as our prior expectations. Rate prior expectations (**Fig S3**) are then provided to the Multinest algorithm to perform nested sampling.

### Multiple hypothesis generation via HypBuilder

Because we perform model selection, we use 5,891 ODE models coded via rule-based modeling in PySB. Each model is generated to include or exclude from 44 reaction rules. There are eight rules that represent division and death for each subtype, and with the potential for three different inter-subtype effects (including none) to have an impact on division or death, each division and death reaction has 3 options, leading to 24 potential rules relating to division/death in total. There are four rules that represent hierarchical phenotypic transitions, which likewise have three potential inter-subtype effects, for 12 rules in total representing hierarchical phenotypic transitions. There are eight rules related to non-hierarchical phenotypic transitions, for 20 total potential phenotypic transition rules out of the 44 rule options.

We use HypBuilder (https://github.com/LoLab-VU/HypBuilder) to automatically generate the 5,891 PySB models that we would otherwise have to code by hand. HypBuilder is software for the automatic generation user-specified collections of mechanistic rule-based models in the PySB format. The input CSV file contains a global list of all possible model components, and reactions, as well as any instructions regarding model creation. The instructions dictate which subsets of model components and reactions will be combinatorially enumerated to create the collection of models. The reactions are parsed via HypBuilder’s molecular interaction library, a library of defined reaction rule sets that is outfitted with common PySB interactions and is customizable to include more interactions should the user need them. Once parsed and enumerated each combination of rules is exported as an executable model via PySB.

The instructions for model construction used in this work direct HypBuilder to use a “list” method to enumerate all candidate models of interest using prior knowledge of likely combinations of model variables (see https://github.com/LoLab-MSM/Bayes-MMI for code used to enumerate candidate models and create the list for HypBuilder).

If the candidate model set contains every relevant biologically plausible possibility, we can consider the entire set of models as representative of 100% of the probability that one of the candidate models explains, or provides the mathematical basis underlying, the data. This is an assumption that cannot truly be met, and most model selection literature acknowledges that one cannot find the “true” model (14,43). However, prior knowledge enables us to determine that all 5,891 models represent all possibilities with regard to outstanding SCLC hypotheses to the best of our ability.

We visualize the prior expectations for the 44 rate parameters as a probabilistic distribution per parameter (prior marginal distribution) (**Fig S3**). Correspondingly, a probabilistic representation of best-fitting rates for each model is returned by the Multinest algorithm (posterior marginal distribution) (e.g., **Fig 4B, 5E**; **Fig S5, S6B**).

### Parameter estimation and evidence calculation by nested sampling

As noted in Eqs. (1) and (2), rate parameters must be set in order to run simulations of a mathematical model. Parameter estimation is the process of determining optimal rates that result in a model simulation recapitulating the data it is meant to represent. Multiple methods exist for parameter fitting or model optimization, (66,67) with Bayesian methods utilizing a prior rate parameter distribution, *P(****θ***), where ***θ*** represents the set of *n* parameters {*θ*_*1*_, *θ*_*2*_, *…, θ*_*n*_}, and a likelihood function to assess a parameter set

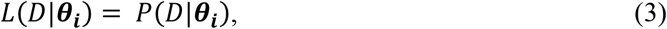

Where ***θ***_***i***_ is the i^th^ parameter set and *D* represents the data being used for fitting. Parameter set ***θ***_***i***_ is scored via the likelihood function *L(D*|***θ***_***i***_) and optimization continues, moving toward better-scoring parameter sets until an optimal score is reached.

For our likelihood function, we represent SCLC tumor steady-state proportion probabilistically, generating a Beta distribution (bounded by 0 and 1) to represent the means and standard deviations of sample replicate subtype proportions, accounting for noise in the proportional space. We test *n-1* subtype proportions to ensure independence of each sampled subtype proportion, to result in the probability of *n-1* independent events (tumor subtype proportions) occurring.

With a prior probability, *P(****θ***), and a likelihood (Eq. 3) the posterior probability can be calculated via Bayes’ Theorem,

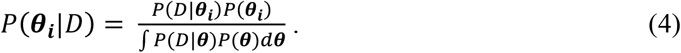

The denominator of Bayes’ Theorem represents the likelihood integrated over all parameter sets, called the marginal likelihood or model evidence. Nested sampling computes this value (Skilling, 2004).

To perform nested sampling, we utilize the Multinest algorithm (46–48). Multinest samples multi-dimensional parameter space, bounding its search by parameter values along each axis in each of the multiple dimensions based on prior expectation of parameters, *P(θ)* input by the user. It removes the lowest-probability parameter set and chooses a new one from within the bounded parameter space, subsequently re-drawing the search space with the bounds incorporating the new parameter set. This continues until all parameter sets representing the bounds of the search space have approximately equal probability, and the algorithm estimates that the remaining probability of parameter sets within the bounds is less than a user-defined tolerance. Each parameter set is evaluated based on a user-defined likelihood function (Eq. 3). Finally, the likelihood values that correspond to each sampled parameter set are arranged in the order they were replaced, and the integral over these is taken to approximate the integral over all possible models, that is, the marginal likelihood or Bayesian model evidence. We used the Multinest-returned importance nested sampling evidence value, because multiple importance nested sampling runs (multiple ‘replicates’) for the same candidate model and prior parameters returned more consistent evidence values than ‘vanilla’ nested sampling (data not shown) (48).

For our likelihood function we represented simulation outcomes – proportions of subtypes at steady state – by a Beta distribution, calculating *α* and *β* using the mean *μ* and variance *σ*^*2*^ of each dataset (68):

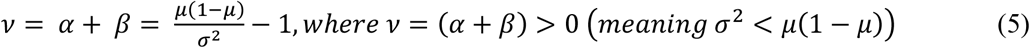

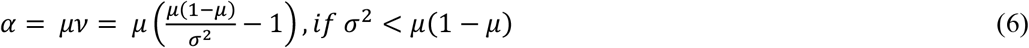

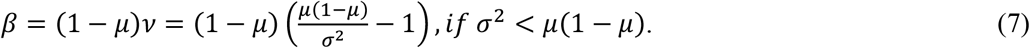

We calculated the log-likelihood of each subtype mean from the dataset being fit against the simulated subtype’s beta distribution,

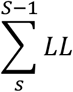

where

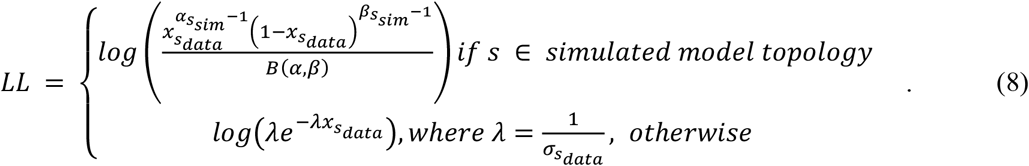

*S* is the set of subtypes, 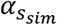 and 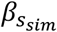 indicate the calculation of *α* (Eq. 6) and *β* (Eq. 7) using the proportion of subtype *s* from the simulation and the variance of subtype *s* in the data, 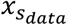 is the mean proportion of subtype *s* in the dataset, *B(α,β)* is 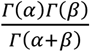 and *Γ* is the Gamma function, and 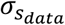 is the standard deviation of the data. Using the exponential function 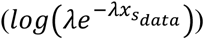 as part of the log-likelihood (Eq. 8) enabled us to calculate a likelihood value for subtypes not present in a model’s topology, which should be a poor log-likelihood if the subtype has a high proportion in the data but was not included in the model topology, or a better log-likelihood if the subtype has a low proportion in the data but was not included in the model topology (and therefore potentially contributing to overfitting). The Python module scipy.stats was used to calculate the Beta log likelihood (Eq. 8, above) and the exponential log likelihood (Eq. 8, below). A simulation would not be scored (return *NaN* and thus be thrown out by the Multinest fitting algorithm) if the tumor subtype proportions did not reach steady state (calculated by whether a proportion timecourse had a slope of zero for the last 7.5% of the simulation).

Multinest is run per model per dataset, which equates to performing 5,891 mechanistic interpretations, 3 times each. CPU time for one model fitting was on average 19 hours (∽0.80 days), with a range of 5 minutes to 28 days. If Multinest had not reached its stopping point by 28 days, we assumed that all regions of parameter space were similarly unlikely and that further running of the algorithm would only continue to refine the search of the unlikely space; models with this difficulty are very likely to have low marginal likelihood due to the unlikeliness of the parameter space. We do not include these incompletely-searched models in our multimodel inference analyses (**Figs 3-5**) and we confirmed that all models that reached 28 days of CPU time without reaching the Multinest stopping point have an extremely low evidence value at the time they were terminated.

### Candidate model prior and posterior probabilities and confidence interval calculation

Each candidate model is considered equally likely prior to fitting by Multinest. That is, every candidate model has an equal prior probability of being the optimal model to represent the underlying SCLC tumor system,

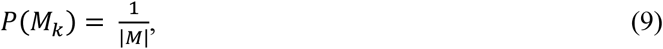

Where *M* is the set of all candidate models. With the model evidence, or marginal likelihood, *P(M*_*k*_) estimated by Multinest, (46–48) the posterior probability per model can be calculated as

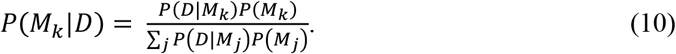

With a posterior probability per model, (**Fig 3**) we calculate a 95% confidence interval. This is accomplished by summing decreasing model posterior probabilities until the sum is 0.95, then considering those models as our 95% CI (14) (**Fig 3**, orange). Using this confidence interval results in ∽1000 models per dataset, a considerable decrease from the initial 5,891. This is a more traditional approach to determining a confidence set of models.

We also took an approach discussed in (14). In this approach, a CI is informed by use of the Bayes Factor between the highest-scoring model and consecutively decreasing scoring models, until the Bayes Factor is larger than a particular cutoff. The models in this CI would be those models *i* for which 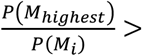 *cutoff*. Burnham and Anderson denote such a method as a “relative likelihood confidence interval” and discuss its support by statistical theory, noting that it is uncommonly found in the model selection literature. We used a cutoff of 10^1/2^, the lowest Bayes Factor at which a difference may be determined (49). Even with this permissive cutoff, the relative likelihood CI includes only tens of models, an even greater decrease from the initial number of candidates.

### Prior and posterior probabilities per hypothesis being investigated

Each hypothesis has an assigned prior probability based on our prior expectations. For all hypotheses, we took an approach where we considered each hypothesis as equally likely compared to competing hypotheses. For the inclusion of most model terms, this was a prior probability of 50%, where it is 50% likely the model term is part of a model that is the best representation of the tumor system, and 50% likely that same term is not part of that model. For the inclusion of effects in the candidate models, the prior probability for a given effect is 33%, where it is equally likely that an effect is generated by Y, generated by A2 and Y, or that no effect is present. The comparison between effect types (including none) is included in **Tables S6-S8**, while the comparison of any effect at all *vs*. no effect (50% *vs*. 50%) is included in the main text.

For the model topology analysis, we considered it equally likely that any model topology could best represent the tumor system that generated each dataset, and with 11 possible model topologies this resulted in a 9% prior probability per model topology (**Fig 4A**). For model initiating subtype hypotheses, (**Fig S2G**) with 15 potential combinations of initiating subtypes, each initiating subtype combination has a 6.67% prior probability.

Each candidate model can then be assigned a prior probability conditional on the hypothesis being considered, *P(M*_*k*_|*H*_*i*_), where *M*_*k*_ is the *k*^*th*^ candidate model and *H*_*i*_ is the hypothesis being considered. The calculation of *P(M*_*k*_|*H*_*i*_) is based on the number of candidate models that fall under the hypothesis being considered,

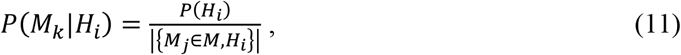

where {*M*_*j*_ *⊂ M, H*_*i*_} is the set of all models assigned to *H*_*i*_. For example, if *H*_*i*_ is the hypothesis that the model term “A to Y transition” is part of the model that would best represent the SCLC tumor system, then {*M*_*j*_ *⊂ M, H*_*i*_} is the set of all candidate models that include the “A to Y transition” model term.

Using this prior probability, the posterior probability for an individual model, conditional on the hypothesis being considered, can be calculated as

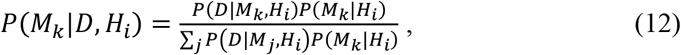

Where *P(D*|*M*_*k*_,*H*_*i*_) is the Bayesian model evidence (marginal likelihood) for *Model*_*k*_.

The posterior probability for an individual model *k* under hypothesis *H*_*i*_, *P(M*_*k*_|*D,H*_*i*_), is not directly used, as the posterior probability of *H*_*i*_ itself, *P(H*_*i*_|*D)* is of principal interest. Under Bayes’ Theorem,

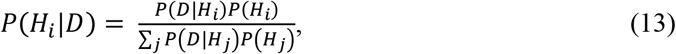

Where *P(D*|*H*_*i*_) is the marginal likelihood of *H*_*i*_ over all models to which it applies, {*M*_*j*_ *⊂ M, H*_*i*_}. According to (49), this can be calculated as

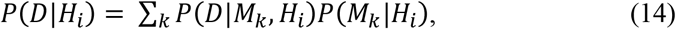

with a summation instead of an integral because each model has a discrete prior probability as calculated in Eq. (11).

Using the results of Eq. (14) in Eq. (13), we then calculate the posterior probability for each hypothesis, pictured in **Fig 5A-D** and noted in **Table 2**. In this way, we can use Bayesian calculation rather than parameter importance analysis (Galipaud et al., 2014, 2017; see Note S1-S3) to determine the posterior probability of each model term. This also enables us to avoid bias in considering models with and without certain model term, if an uneven number of candidate models contain a model term *vs*. do not contain the term (69).

### Posterior odds per hypothesis being investigated

All model terms and variables begin with a prior probability of 0.5. With equal prior probabilities across all model term hypotheses, the posterior odds represented by 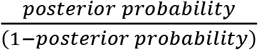 is equivalent to the Bayes Factor. Therefore, calculation of the posterior odds and the Bayes Factors for each model term are equivalent.

A posterior probability of model term inclusion of 0.75 or more, or probability of 0.25 or less, would be considered substantial evidence for inclusion or exclusion of that term, respectively (49). Given the nature of the posterior odds, where a value of 2 indicates that one hypothesis is twice as likely to be true as the other, we also consider posterior probabilities of 0.667 or more, or 0.333 or less, to be notable evidence for inclusion or exclusion of the model term considered. We consider probabilities between 0.333 and 0.667 to not have been significantly informed by the data.

### Bayesian model averaging of parameter sets

Since Multinest returns multiple best-fitting parameter sets, each parameter in a model has a frequency distribution representing the values it takes on over these parameter sets. We thus consider each parameter using a probabilistic representation, per model (posterior marginal distribution) (**Fig 4B, 5E**; **Fig S5, S6B**). Since each candidate model is assigned a posterior probability as in Equation (10), all best-fitting parameter sets for that model can be assigned the same posterior probability. The frequency distribution of one parameter’s values across a model’s best-fitting parameter sets are thus weighted by its model’s posterior probability. Then, the frequency distributions of weighted parameter values per model can be combined, representing the distribution of potential values of a particular parameter, weighted by model posterior probabilities. This way, parameter values in the distribution that come from models with a higher posterior probability (thus higher model evidence) will have more of an effect on the probabilistic representation, since they represent more likely values for the parameter.

To assemble representative fitted parameter sets for each candidate model, we used the first 1000 parameter sets from the Multinest equally weighted posterior samples per model. With up to 44 parameters and up to 5,891 models, the collection has 44 parameter columns and up to 5,891,000 rows representing a parameter vector. The collections were made per dataset.

### Comparing parameter distributions

As above, each kinetic parameter has a frequency distribution representing 1000 fitted values per candidate model, meaning up to 5,891,000 fitted values across all models (weighted using Bayesian model averaging, as above). To compare parameter rates across models in the same dataset but with different topologies, we grouped each parameter according to the model topology from which it came. We then sampled 1000 values from the BMA-weighted distribution per kinetic parameter across all models of the same topology. We performed ANOVA followed by Tukey HSD at family-wise error rate (FWER) of 0.01, using the Python module statsmodels. Below an FWER of 0.01, we considered the sampled parameters significantly different across models. We then repeated the sampling, ANOVA, and Tukey HSD for a total of 10 iterations. We then averaged across determinations of significant/non-significant and if a parameter comparison across model topologies was significantly different more often than it was not different, we considered the parameter rates to be different comparing model topologies. The same methodology was used to compare parameter rates across different datasets.

### Generating a consolidated model of the SCLC tumor

A hypothesis (model term) whose posterior probability is further from its prior probability indicates more information gained during the nested sampling process - more knowledge provided by the data. Conversely, a posterior probability similar to the corresponding prior probability indicates that the data did not inform our prior knowledge.

To unify the varying models into one view of SCLC biology, we brought together model probabilities from each three-subtype topology per dataset (**Fig 5E**). To bring together the results for each three-subtype topology results in the investigation of what appears as a four-subtype topology. In fact, if we are to envision one model that can represent one system that generated all three datasets, it would need to include all four subtypes. We consider this a reasonable practice in that all transition posterior probabilities in the three-topology subtypes either were little informed by the data or had a value indicating that transitions are likely; in addition, all Non-NE effects were either little informed by the data or had a value indicating that these effects are unlikely. Posterior probabilities were not the same between three-subtype topologies, but these trends of likely or unlikely model features generally agreed.

When consolidating models in this way, if model terms were part of multiple topologies (e.g., the A-to-N transition is part of the A, N, and Y topology, best representing the RPM dataset, and the A, N, and A2 topology, best representing the SCLC-A cell line dataset) we took the posterior probability of the model feature closer to 0.5. For example, the posterior probability for the A to N transition in the RPM dataset is 0.709 and the posterior probability for this same transition in the SCLC-A dataset is 0.626. Therefore, in the four-subtype consolidated representation, the posterior probability for the A to N transition is 0.626. This is the most conservative way to represent the knowledge gained by the data from the perspective of the entire SCLC system, allowing for the most uncertainty to remain. We consider this practice as avoiding claiming more certainty about model features than the data may provide.

## CONTRIBUTIONS

Conceptualization, S.P.B, L.A.H., and C.F.L; methodology, S.P.B., L.A.H., and M.A.K.; software, S.P.B., L.A.H., and M.A.K.; formal analysis and investigation, S.P.B.; resources, C.F.L., J.S. and V.Q.; data curation, S.P.B.; writing – original draft, S.P.B., C.F.L.; writing – review & editing, S.P.B., L.A.H., M.A.K., J.S., V.Q., C.F.L.; visualization, S.P.B.; supervision, V.Q., L.A.H., and C.F.L; funding acquisition, V.Q. and C.F.L.

## ACKNOWLEDGMENTS

The authors would like to thank Sarah Groves, Michael Irvin, and Christine Lovly for insightful conversations and critical feedback on this work. This work was supported by the following funding sources: S.P.B. was supported by the National Institutes of Health (NIH) [T32GM007347 and T32LM012412] and the National Cancer Institute (NCI) [F30CA247078]; C.F.L. was supported by the National Science Foundation (NSF) [MCB 1411482] and NSF CAREER Award [MCB 1942255]; C.F.L. and V.Q. were supported by the National Institutes of Health (NIH) [U54-CA217450 and U01-CA215845].

## AVAILABILITY

All the code to reproduce the data analysis and the figures is open source and can be found in this GitHub repository: https://github.com/LoLab-VU/Bayes-MMI. This repository contains all the source code for each step in the analysis.

Files needed to reproduce the figures without running the data analysis prior can be found in this DropBox folder: https://www.dropbox.com/sh/4fqzpvu9hgyjicm/AABdfFlCenEuiOPgiH0TT-xqa?dl=0. Download of these files and placement in the corresponding directories in the GitHub repository enables reproduction of Figures 3-5.

## Notes

https://github.com/LoLab-VU/Bayes-MMI

https://www.dropbox.com/sh/4fqzpvu9hgyjicm/AABdfFlCenEuiOPgiH0TT-xqa?dl=0

### Competing Interest Statement

The authors have declared no competing interest.

### Summary of Updates

Increasing figure resolution.

