## Supplemental Information for "Unified Tumor Growth Mechanisms from Multimodel Inference and Dataset Integration"

### Note S1. A didactic example to contrast AIC vs posterior probability calculated by Bayes-MMI for model selection and multi-model inference

#### 1.1 Using multiple models to evaluate how well a variable informs the observed data: an example

Mathematical models are a useful way to interrogate a biological system. Building a model can help the investigator hypothesize relationships within the system, and more specifically evaluate whether and how these relationships relate to observed phenomena. Each piece of a model represents a distinct hypothesis; thus the model as a whole represents multiple hypotheses about the system of interest.

For example, a farmer might wish to determine which aspects of his farm most affect his monthly income: milk from his cows, eggs from his hens, wool from his sheep, or pony rides offered to the public. He is interested in assessing how he should price the product or service based on how many animals he must keep his income steady. Additionally, if a provided product or service doesn't impact his monthly income, perhaps he will stop providing it. This is a question with one outcome or response variable, the monthly income, and up to four predictor variables, one each for the animals that provide the products or services. We can write a linear equation that describes these interactions as shown below:

$$income = p_{milk} * cows + p_{eggs} * hens + p_{wool} * sheep + p_{rides} * ponies + savings\ interest \quad (1)$$

$$y = \beta_1 x_1 + \beta_2 x_2 + \beta_3 x_3 + \beta_4 x_4 + \beta_0 \quad (2)$$

(Eq. 2) represents a linear regression model corresponding to the more intuitively written (Eq. 1), where the savings interest represents the "intercept", or a baseline amount of money where the farm budget starts each month. However, (Eq. 2) is only one of many possible models that the farmer could use to determine how these variables come together to yield his income and perhaps improve on his earnings. A multi-model averaging approach would enable the farmer to explore model hypotheses to explore how much he should charge, $\beta_{1-4}$, based on number of working animals that month, $x_{1-4}$, and his monthly income, $y$, *and,* how much each of those animals $x_{1-4}$ contribute to the income $y$.

Besides using linear regression to determine pricing, the farmer can evaluate all potential models including few, some, or all farm animals to see which animals most directly affect his income. Perhaps the lower-earning products per month are not needed to maintain the income based on the numbers of animals providing each product. For example, if sheep's wool and pony rides do not bring in as much income as milk or eggs, maybe they do not contribute meaningfully to the income, and the best model would look like this:

$$income = price_{milk} * cows + price_{eggs} * hens + savings\ interest \quad (3)$$

$$y = \beta_1 x_1 + \beta_2 x_2 + \beta_0 \quad (4)$$

With all possible combinations of animals that the farmer might use to support his income, there are 16 different possible models that can help predict his monthly income, below:

$$y = \beta_0$$

$$y = \beta_1 x_1 + \beta_0$$

$$y = \beta_2 x_2 + \beta_0$$

$$y = \beta_3 x_3 + \beta_0$$

$$y = \beta_4 x_4 + \beta_0$$

$$y = \beta_1 x_1 + \beta_2 x_2 + \beta_0$$

$$y = \beta_1 x_1 + \beta_3 x_3 + \beta_0$$

$$y = \beta_1 x_1 + \beta_4 x_4 + \beta_0$$

$$y = \beta_2 x_2 + \beta_3 x_3 + \beta_0$$

$$y = \beta_2 x_2 + \beta_4 x_4 + \beta_0$$

$$y = \beta_3 x_3 + \beta_4 x_4 + \beta_0$$

$$y = \beta_1 x_1 + \beta_2 x_2 + \beta_3 x_3 + \beta_0$$

$$y = \beta_1 x_1 + \beta_2 x_2 + \beta_4 x_4 + \beta_0$$

$$y = \beta_1 x_1 + \beta_3 x_3 + \beta_4 x_4 + \beta_0$$

$$y = \beta_1 x_1 + \beta_2 x_2 + \beta_3 x_3 + \beta_4 x_4 + \beta_0$$

The model selection process that the farmer can use involves evaluating how well each candidate model's (from this superset of plausible models) simulations of *pricing * animal* compared to the observations of his income. This process is often performed for choosing one best-matching model, and whichever model is best is often considered to include only the important model variables (the most important animals from the perspective of the farmer's income. However, we and others argue, one may use model averaging, where parameter values (prices) can be weighted by model probability and then combined into a distribution of likely values, and similarly variable (animal) likelihood can be weighted and summed across all models (1,2). Thus, this set of candidate models can provide support for or against pieces of models, and therefore support for or against hypotheses.

What is the best framework for which the farmer can perform model selection and model averaging? We argue that a Bayesian framework is optimal, compared in this example to the most common model selection/model averaging framework, the information theoretic approach of AIC (1,3). In sections 1.2-1.6 we provide the background information that motivate our claim that a Bayesian framework should be optimal. In section 1.7, we provide our Bayesian analysis of a published linear regression example that investigated AIC model selection and model averaging to compare the two frameworks.

### 1.2 Marginal likelihood or "evidence" is calculated using model optimization followed by Bayes' Theorem

In a Bayesian statistics context, probability indicates the degree of belief or confidence in an event, or in a proposal that a statement may be true – that is, a hypothesis. This prior degree of belief (hypothesis) is then updated with new data, which can yield evidence that our prior belief must be modified. In this case, data can refer to measurements related to the hypothesis and is synthesized into a likelihood that the hypothesis is true. Prior knowledge combined with data leads to the posterior, updated, probability, defined using Bayes' Theorem as

$$P(H|D) = \frac{P(D|H)P(H)}{P(D)} = \frac{P(D|H)P(H)}{\int P(D|H)P(H)dH} \quad (5)$$

where $P(H|D)$ denotes a conditional posterior probability of the hypothesis ($H$) given the data $D$, $P(D|H)$ denotes the probability of observing the data $D$ if the hypothesis $H$ were true (also called the likelihood), and $P(H)$ indicates prior probability (degree of belief) of the hypothesis before being presented with more data. Calculation of the posterior probability of the hypothesis $P(H|D)$ requires dividing the likelihood of one hypothesis times its prior, $P(D|H)P(H)$, by $P(D)$ (Eq. 5, middle). When comparing many models,

$P(D)$ becomes the integral over the likelihood times the prior probability for each hypothesis $H$ within the set of all hypotheses $\mathcal{H}$ (Eq. 5, right).

When the hypothesis $H$ in (Eq. 5) represents a candidate model $M$ with $n$ parameters, we can think of model $M$ as the set of parameters, $\boldsymbol{\theta_M} = \{\theta_1, \theta_2, \ldots, \theta_n\}$. Each parameter $\theta_i$ itself represents one dimension of the model's probability space. Model optimization to data D assigns a likelihood to each $\boldsymbol{\theta_M}$ in probability space, and these values over the entire parameter space represent the numerator in (Eq. 5) with $H = \boldsymbol{\theta_M}$, $P(D|\boldsymbol{\theta}_M)P(\boldsymbol{\theta}_M)$, a distribution proportional to the probability distribution of model $M$. To calculate the probability distribution of $M$, we must also calculate the denominator of (Eq. 5), $\int P(D|\boldsymbol{\theta}_M)P(\boldsymbol{\theta}_M)d\boldsymbol{\theta}$, the so-called marginal likelihood or Bayesian evidence that normalizes the distribution of model $M$ over all $n$ parameters.

The marginal likelihood is often considered the average of the likelihood over the prior space (4–6). Consider a multidimensional parameter space for a model $M$ where each point in space has an assigned likelihood. Then, if we add another parameter – another dimension – that is only mildly informative, the likelihood values barely change, and they are now spread across even more space. Thus, less evidence is present in the overall space, and therefore adding a parameter to a model will result in a lower marginal likelihood, unless that parameter allows the model $M$ to fit the data $D$ significantly better. In this way, the marginal likelihood intrinsically penalizes more complex models, unless a more complex model better matches the data (4–6).

#### 1.3 AIC is calculated as an estimate of the Kullback-Liebler divergence

The calculation of an information criterion is based on estimating the difference between candidate models and the "true" model (reality). The difference between statistical models or probabilities is measured by Kullback-Leibler (KL) divergence, represented by

$$I(f,g) = \int f(x)\ln\left(\frac{f(x)}{g(x|\boldsymbol{\theta})}\right)dx \qquad (6)$$

where *f* and *g* represent probability distributions. In a typical KL divergence calculation, *f(x)* is the reference distribution, while *g(x)* is the distribution being compared. When using KL divergence in information theoretic model selection, as in (Eq. 6), we consider *f(x)* to be "reality," while $g(x|\boldsymbol{\theta})$ is a candidate model parameterized by $\boldsymbol{\theta}$, which are estimated from data *D* (as above). Here, $g(x|\boldsymbol{\theta})$ is a probability distribution of a model akin to the probability distribution of model *M*, $P(\boldsymbol{\theta_M}|D)$ noted above. *I* represents information, and because *g* cannot exactly match reality, *I(f,g)* represents the information lost when approximating reality *f* by the model *g* (3).

The Akaike Information Criterion, or AIC, is an estimate of this KL divergence based on the likelihood function at its maximum point (1). It also considers the number of parameters in the model $g(x|\boldsymbol{\theta})$, which reduces bias in the estimate of the KL divergence for that model. Therefore, the AIC is denoted by

$$\text{AIC} = -2\ln\left(L(\boldsymbol{\theta}_{best}|D)\right) + 2K = -2\ln\left(P(D|\boldsymbol{\theta}_{best})\right) + 2n \qquad (7)$$

where $L(\boldsymbol{\theta}_{best}|D)$ is the likelihood of the best-fitting parameter set and *K* is the number of parameters in the model (noted as $P(D|\boldsymbol{\theta}_{best})$ and *n*, respectively, to correspond with the Bayesian definitions in 1.2).

#### 1.4 Notable differences between AIC and Bayesian evidence / posterior probability

While they both aim to rank models from a set of candidates so that one or few can be chosen as the best models, there are several key differences between Bayesian evidence/posterior probability and AIC.

As noted in 1.2, the Bayesian evidence or marginal likelihood is represented by $\int P(D|\boldsymbol{\theta}_M)P(\boldsymbol{\theta}_M)d\boldsymbol{\theta}$. Since model optimization to data *D* assigns a likelihood to each value over the entire parameter space, and is represented by $P(D|\boldsymbol{\theta}_M)P(\boldsymbol{\theta}_M)$, integrating over this thereby integrates over every likelihood value across all evaluated parameter values in each dimension. In this way, the marginal likelihood takes into account every parameter set evaluated during optimization. This represents a more comprehensive score for a model compared to AIC, which as in (Eq. 7) only considers the best-fitting parameter set $\boldsymbol{\theta}_{best}$. Additionally, as noted with an example in 1.2, the marginal likelihood penalizes for model complexity (number of parameters). AIC also includes the number of parameters $K$ in its calculation as seen in (Eq. 7); however, this is again a less comprehensive means of penalizing for number of parameters compared to marginal likelihood.

We then consider the inclusion of prior expectations. At the level of model optimization, both AIC and marginal likelihood can be said to include prior expectations (depending on the algorithm used for optimization). Marginal likelihood includes prior expectations by default, represented by the $P(\boldsymbol{\theta}_M)$ term in the integral above. AIC requires only the best-fitting parameter set, so if optimization was performed via Bayesian principles where prior expectations for parameters were assigned, then these priors could well be said to affect $\boldsymbol{\theta}_{best}$ in the AIC calculation. However, Bayesian principles enable inclusion of prior expectations at the level of model ranking. Often, candidate models are considered equally likely a priori, (4–6) and will thus have a prior probability of $\frac{1}{n}$. When considering H in (Eq. 5) equal to a particular model $M_i$, the posterior probability $P(M_i|D)$ firstly *requires* a prior probability $P(M_i)$ for its calculation, and once calculated can be compared to that prior probability $\frac{1}{n}$. In this way, Bayesian principles provide a numerical metric to assess how data informed our knowledge about $M_i$. This plays a role in model averaging as well (see 1.6). AIC does not have a formal means for including prior probability in its calculation nor in the generation of a posterior probability. Probability in the use of AIC is an interpretation of relative likelihoods across candidate models and is not a true probability (see Eqs. 4 and 5 in the next section).

Given the incorporation of the full parameter space and prior expectations, as well as the ability to calculate a Bayesian probability that provides insight into knowledge gained from the data, we consider marginal likelihood the optimal means for model ranking and interpretation therein. A specific model selection example where AIC and marginal likelihood/posterior probabilities are compared is detailed in section 1.7.

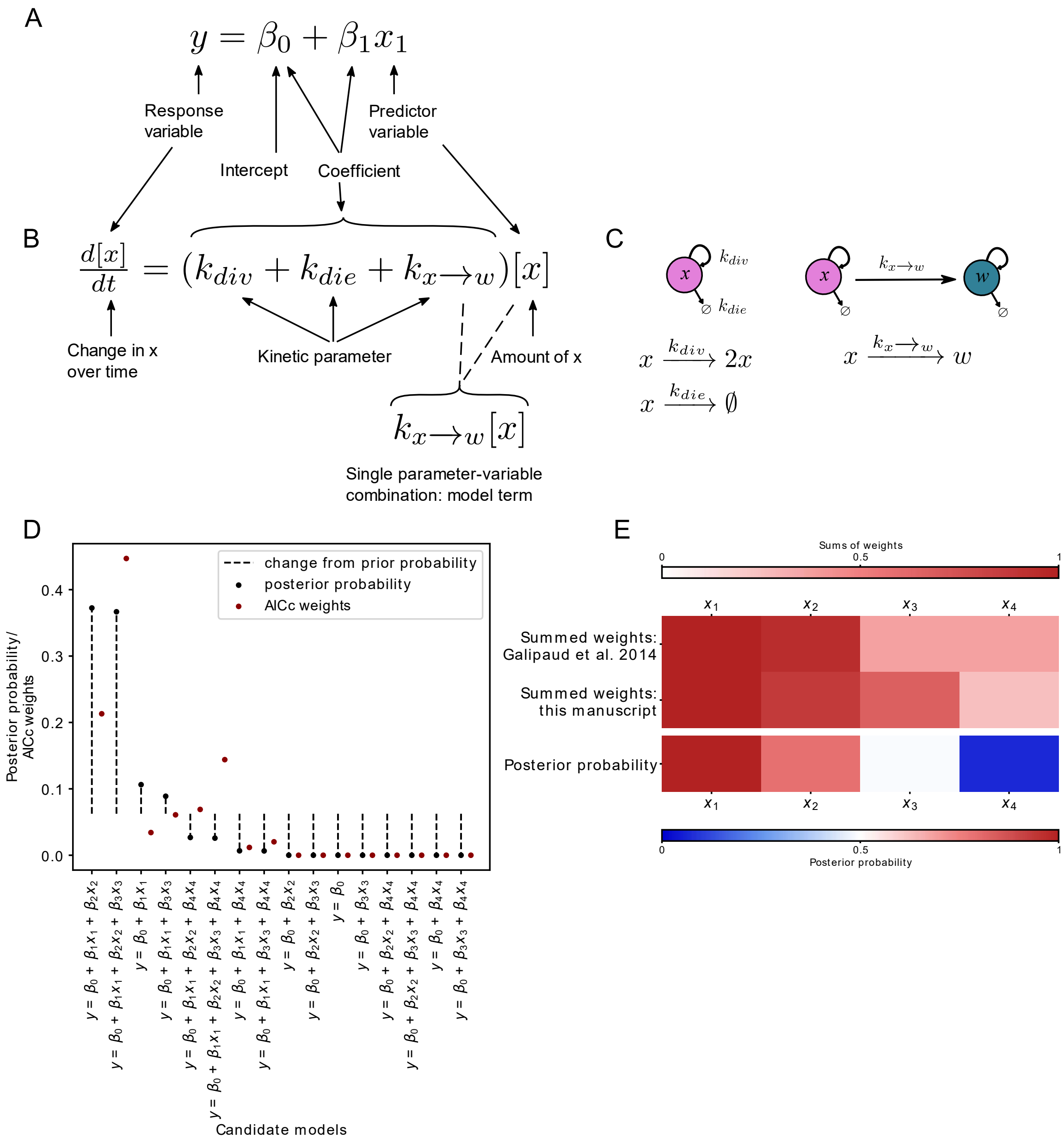


**Fig S1. Bayesian inference better assesses parameter inclusion in the "true" model compared to Akaike Information Criterion.** (A) Aspects of linear regression model assessed by model selection and model averaging. (B) Aspects of mass-action kinetics model / ordinary differential equation assessed by model selection and model averaging. (C) Schematic representation of the equation in (B). (D) Results from our analysis of candidate models using candidate set and data generation code from Galipaud et al., 2014 [10]. Candidate models are arranged along the x-axis by posterior probability. Each posterior probability from our analysis is compared to the AICc weight from our analysis. (E) Heatmap for each model averaging analysis based on data generation and candidate models in [10]. Top two rows include summed weights (SW) published in the original Galipaud et al., 2014 [10] manuscript and from our analysis. Top color bar legend corresponds to these two rows, where SW is a value starting at 0 and weights are summed until the final SW value. Bottom row is the posterior probability calculation from our analysis of this data and candidate set. Bottom color bar legend corresponds to this row, where color represents the probability of the variable's inclusion in the "true" model. Since prior probability starts at 0.5 (white), deeper colors indicate a larger deviation from the prior, with red vs blue indicating more likely or less likely probability, respectively.

#### 1.5 Model selection allows us to evaluate which variables or terms have the largest effect on observed data

Model selection is the process of investigating candidate models, each of which may fit the data to a different extent, while prioritizing model simplicity (aka *lex parsimoniae* (7)). That is, the best model must comprise a balance between matching the data exactly and having the fewest variables and parameters required to do so. In linear regression, such as in the following example, this involves investigating which predictor variables with their fitted coefficients, best match the response variable (**Fig S1A**). In kinetic models, such as those with which we aim to capture the behavior of small cell lung cancer (SCLC), this involves investigating which variables and which model terms, with their fitted kinetic parameters, best match the data (**Fig S1B,C**).

For Bayesian model selection, the hypotheses $H$ in (Eq. 5) represent candidate models $M$ within the set of all models $\mathscr{M}$, and data $D$ represents the dataset to which each model is fitted. Then, the likelihood of that model $M$, $P(D|M)$, is the marginal likelihood. Calculating the marginal likelihood for all candidate models and normalizing by the sum of this value for all models results in the probability of the model $M$ given the data, $P(M|D)$. The higher the probability, the more likely the model best represents the process that generated the data (**Fig S1D**, black points).

Models can be ranked by AIC value alone, and in this case a lower AIC value indicates that a model is a better representation of the data. However, those using AIC can scale the models with respect to the minimum AIC value (1,3). This results in AIC differences,

$$\Delta_i = AIC_i - \min{(AIC)} \qquad (8)$$

where *min(AIC)* is the AIC value of the lowest-scoring (best) model candidate. Using AIC differences, Akaike weights are calculated,

$$w_i = \frac{e^{-\Delta_i/2}}{\sum_{r=1}^{R} e^{-\Delta_i/2}} \qquad (9)$$

where $R$ is the number of models in the candidate set. Akaike weights represent relative likelihoods of the models given the data, and are interpreted as probabilities (1). From this perspective, the higher the Akaike weight, the more likely that model best represents the process that generated the data (**Fig S1D**, red points).

#### 1.6 Model averaging uses model selection outcomes from all models to demonstrate how the observed data informed the model variables or terms that represent our hypotheses

Probabilities or weights are then used for model averaging. Model averaging is a process used when no clear single best model can be identified after model selection. Model averaging can reveal a model variable or term common to better-performing models, indicating that that the process represented by this variable or term is likely to play a role in the process that generated the data, and allowing the user to identify the parts of a model that best capture the data. In our application, this approach implies a departure from a deterministic single model to a probabilistic understanding of mechanisms, with the probabilities determined from the available data.

In Bayesian model averaging, (BMA) each model is weighted by its marginal likelihood or posterior probability (8) and the likelihood of model with particular term or variable can be used to generate an overall likelihood for that term or variable. Here, the prior probability of a term, which we typically consider to be equally likely vs. unlikely before model optimization, is important. Now, the $H$ of (Eq. 5) indicates the hypothesis that a particular model term (individual process within the system of interest) is part of the “true” model, and the prior $P(H)$ and likelihood of that term $P(D|H)$ can be used to calculate $P(H|D)$. Thus we can investigate how the data informed our knowledge about $H$, from its prior to posterior probability.

To determine whether to predict that a model variable or term is likely to be involved in the system of interest, the Bayes Factor can be used. The Bayes Factor is the ratio of the likelihood of two hypotheses, and when prior probabilities are equal is equivalent to the ratio of the two posterior probabilities:

$$BF = \frac{P(D|H_1)}{P(D|H_2)} = \frac{P(H_1|D)/_{P(H_1)}}{P(H_2|D)/_{P(H_2)}}; BF = \frac{P(H_1|D)}{P(H_2|D)}\ when\ P(H_1) = P(H_2) \quad (10)$$

Here, $H_1$ may represent the hypothesis that the model variable does play an important role, while $H_2$ may represent the hypothesis that it does not. The Bayes Factor (BF) between one hypothesis and another can be considered a true difference between the probability of each hypothesis when $10^{-1/2}$ > BF > $10^{1/2}$. The value of $10^{1/2}$ (~=3) is the lowest at which a difference may be determined; or $10^{-1/2}$ (~= ⅓) when the higher-probability hypothesis is in the denominator of the BF (9). In this way, comparing hypotheses via BF can result in one of three outcomes: $P(H_1|D)$ is > 3 times more likely than $P(H_2|D)$, meaning hypothesis $H_1$ is informed by the data to be significantly more likely than $H_2$; $P(H_1|D)$ is < ⅓ times less likely than $P(H_2|D)$, meaning $H_1$ is informed by the data to be significantly less likely than $H_2$; or, $P(H_1|D)$ is between ⅓ less likely and 3 times more likely than $P(H_2|D)$, meaning $H_1$ is not informed by the data to be significantly different than $H_2$. We simply denote this last outcome as "$H_1$ is not informed by the data."

When using information criteria such as AIC, parameter importance analysis is typically performed to investigate how well the data supports a model variable. For each variable, the Akaike weights of all models containing that variable are summed (1). The sum of weights (SW) is treated as the probability that a variable belongs in the "true" model, thus the biological feature represented by this variable plays an important role in the biological system. There does not appear to be an accepted threshold over which the SW for a variable indicates it should be included in a model, thus it seems practitioners choose a threshold on an ad hoc basis (10).

### 1.7 Advantages of Bayes-MMI over AIC for model selection and model averaging: continuing example

Despite equivalent goals between BMA and SW, problems have been noted with the latter. To illustrate this point, we consider the example of (10), and compare AIC-based SW to our Bayesian method combining marginal likelihood results of model selection and BMA of terms across candidate models, named Bayes-MMI.

In (10), the investigators generated "ground truth" data using four variables $x_{1\text{-}4}$, assigning differing correlations with the response variable $y$, including no correlation between $x_4$ and $y$ (Pearson correlation coefficient 0.0) (**Note S2**). A variable with no relation to the response variable should not appear in high-scoring models, and should receive a low SW. Galipaud and colleagues performed model selection using all variable combinations including intercept-only, (16 models) calculated AICc, (AIC corrected for small sample sizes) and assessed parameter importance using SW. They found that while $x_4$ was completely unrelated to the response, it appeared among best-ranked models and had a SW different from zero (10).

To test how Bayes-MMI compares to AIC-based SW methods, we used data simulated via the code provided in (10), we ran a nested sampling analysis via PyMultinest (4–6) (see **Methods**) using the same 16 candidate models, and obtained similar results to Galipaud and colleagues (**Table S1**; **Table S3**; **Note S2**). Since nested sampling yields the marginal likelihood, (4–6), we can leverage Bayesian principles as noted above to explore model hypotheses space and identify the best candidate models. We use candidate model posterior probabilities to perform our own ranking of models (**Fig S1D**; **Table S2**). We also calculate variable posterior probabilities, (**Fig S1E; Table S3**) using equivalent prior probabilities of variable inclusion: prior probability that the inclusion of a variable in the "true" model is 0.5, or completely unknown (equivalent probability that it is present vs that it is not present).

Firstly, while according to AICc, $x_4$ can be found in the best-ranked models, when ranking according to posterior probability, $x_4$ does not appear until the fifth-highest ranked model (**Fig S1D; Table S1, S2**;(10)).

The four highest-ranked models have a cumulative probability of 0.934 (**Fig S1D; Table S2**). Thus, most of the probability that one candidate model is the best model is contained in those models, which more accurately do not include $x_4$. In fact, $x_4$ appears only in candidate models whose posterior probability has *decreased* compared to the prior probability (**Fig S1D,** dotted lines above *vs* below posterior probability points; **Table S2**). Next, while the SW for $x_4$ in our AICc analysis is 0.25, the posterior probability that the inclusion of $x_4$ is supported by the data is 0.07 (**Table S3**).

Using Bayesian principles, we consider the prior probability of $x_4$ inclusion equivalent between presence and absence (probability of 0.5), but after incorporating data, the posterior probability of $x_4$ inclusion has decreased to a large extent, to a probability of 0.07. We view this as a reasonably low probability, given that $x_4$ is not at all correlated to $y$. If $H_1$ represents "$x_4$ should be included in the model", and $H_2$ the opposite, the BF for $H_1$ vs $H_2$ is 0.07/(1-0.07) = .075 < ⅓, and we determine that $x_4$ should not be included in the model. The case of $x_3$ is also instructive, where the prior probability of inclusion at 0.5 has barely changed to a posterior probability of 0.49 after incorporation of data (BF = .49/(1-.49) = 0.96). With the weak correlation of $x_3$ to $y$ at 0.05, Bayesian inference has indicated that, at least with the data used, a definitive choice to include or exclude $x_3$ cannot be made. The SW for $x_3$ is 0.67, and as such a practitioner would likely include it in a chosen model, possibly without considering that such a weakly correlated variable could achieve this SW.

Therefore, we consider using Bayesian analysis based on the marginal likelihood (Bayesian evidence) to be a superior way to perform model selection, and specifically analysis of variable or model term inclusion or exclusion. We thus aim to employ multimodel inference (MMI; model selection and model averaging together) in the context of Bayesian statistics. There have been biological investigations using MMI approaches (11,12) but, to our knowledge, Bayesian MMI has not previously been applied to our models of interest, cell population dynamics models. Using MMI to assess the posterior probability of different model variables is comparable to Bayesian variable selection, which in biomedicine has been used to determine genetic loci associated with health and disease outcomes in linear models (12). We find using a

Bayesian approach to MMI results in probabilities that biologically relevant model features are (or are not) supported by the data, and is likely relevant to any cancer or developmental biology application and can be used to investigate model variables even in the context of limited or uncertain data.

### Note S2. Comparing Galipaud et al., 2014 to our own analysis of the same simulated data

In (10), the investigators generated "ground truth" data using four variables $x_{1\text{-}4}$ with differing correlations with the response variable *y*. $x_1$ was strongly correlated to response variable *y* with Pearson correlation coefficient $r_{y,x_1}$ of 0.70, $x_2$ moderately correlated with $r_{y,x_2}$ 0.20, $x_3$ weakly correlated with $r_{y,x_3}$ 0.05, and $x_4$ uncorrelated with $r_{y,x_4}$ = 0.0. They performed model selection using variable combinations including intercept-only, (16 models) calculating AICc (AIC corrected for small sample sizes) and assessed parameter importance using SW (10). Galipaud et al. used least squares regression via the R lm() function, while for our nested sampling parameter search we used a least squares likelihood function with PyMultinest searching the 5-dimensional parameter space ($x_1$, $x_2$, $x_3$, $x_4$, and $\beta_0$) using uniform prior distributions of 0 to 10 for each response variable and -10 to 10 for the intercept $\beta_0$. Our nested sampling results are similar to those in (10), as seen in **Table S1** and **Table S3** columns "SW: Galipaud" and "SW: this manuscript".

**Table S1. Summary of nested sampling model selection results on the simulated dataset and model selection problem in Galipaud et al., 2014.**

| $\beta_0$ | $x_1$ | $x_2$ | $x_3$ | $x_4$ | *k* | rank, AICc | rank (AICc) in (10) | *log(L)* | AICc | $\Delta_i$ | $w_i$ |
|---|---|---|---|---|---|---|---|---|---|---|---|
| -4.942 | 0.661 | 0.197 | 0.13 | | 5 | 0 | 1 | -45.699 | 99.82 | 0 | 0.447 |
| -3.481 | 0.651 | 0.192 | | | 4 | 1 | 0 | -47.526 | 101.301 | 1.482 | 0.213 |
| -4.753 | 0.657 | 0.19 | 0.118 | 0.005 | 6 | 2 | 3 | -45.724 | 102.085 | 2.266 | 0.144 |
| -3.806 | 0.666 | 0.207 | | 0.002 | 5 | 3 | 2 | -47.568 | 103.556 | 3.737 | 0.069 |
| -2.881 | 0.668 | | 0.117 | | 4 | 4 | 5 | -48.778 | 103.807 | 3.987 | 0.061 |
| -1.66 | 0.662 | | | | 3 | 5 | 4 | -50.419 | 104.962 | 5.142 | 0.034 |
| -3.009 | 0.674 | | 0.124 | 0 | 5 | 6 | 7 | -48.794 | 106.009 | 6.189 | 0.02 |
| -1.658 | 0.661 | | | 0.001 | 4 | 7 | 6 | -50.424 | 107.098 | 7.278 | 0.012 |
| 1.664 | | 0.222 | 0.109 | | 4 | 8 | 9 | -95.065 | 196.38 | 96.56 | 0 |
| 2.811 | | 0.215 | | | 3 | 9 | 8 | -96.334 | 196.791 | 96.971 | 0 |
| 1.601 | | 0.23 | 0.105 | 0 | 5 | 10 | 13 | -95.081 | 198.582 | 98.762 | 0 |
| 2.784 | | 0.217 | | 0 | 4 | 11 | 11 | -96.336 | 198.921 | 99.102 | 0 |
| 3.994 | | | 0.099 | | 3 | 12 | 12 | -98.919 | 201.961 | 102.142 | 0 |
| 4.973 | | | | | 2 | 13 | 10 | -100 | 202.041 | 102.221 | 0 |
| 4.021 | | | 0.095 | 0.001 | 4 | 14 | 15 | -98.931 | 204.112 | 104.292 | 0 |
| 4.973 | | | | 0 | 3 | 15 | 14 | -100.001 | 204.126 | 104.306 | 0 |

Candidate models are ranked by AICc as in Galipaud et al., 2014; rank in our analysis can be compared to the ranking in Galipaud et al., 2014 ("rank" *vs*. "rank in Galipaud et al., 2014" columns). Maximum log-likelihood parameter estimates and AICc are calculated from PyMultinest output. Parameter estimates are reported if present for each of the 16 candidate models. *k*, total number of estimable parameters; *log(L)*, maximum log-likelihood returned by our likelihood function; AICc, AIC "corrected" for small sample size; $\Delta_i$, *AICc – min(AICc)* per model; $w_i$, Akaike weight.

**Table S2. Summary of nested sampling model selection results on the simulated dataset and model selection problem in Galipaud et al., 2014.**

| ***$\beta_0$*** | ***$x_1$*** | ***$x_2$*** | ***$x_3$*** | ***$x_4$*** | ***k*** | **rank, post. prob.** | **rank, AICc** | **rank (AICc) in Galipaud et al. 2014** | ***log(Z)*** | ***log(Z)* error (+/-)** | **prior prob.** | **post. prob.** |
|---|---|---|---|---|---|---|---|---|---|---|---|---|
| -3.481 | 0.651 | 0.192 | | | 4 | 0 | 1 | 0 | -55.669 | 0.047 | 0.062 | 0.372 |
| -4.942 | 0.661 | 0.197 | 0.13 | | 5 | 1 | 0 | 1 | -55.685 | 0.052 | 0.062 | 0.367 |
| -1.66 | 0.662 | | | | 3 | 2 | 5 | 4 | -56.923 | 0.043 | 0.062 | 0.106 |
| -2.881 | 0.668 | | 0.117 | | 4 | 3 | 4 | 5 | -57.102 | 0.048 | 0.062 | 0.089 |
| -3.806 | 0.666 | 0.207 | | 0.002 | 5 | 4 | 3 | 2 | -58.302 | 0.054 | 0.062 | 0.027 |
| -4.753 | 0.657 | 0.19 | 0.118 | 0.005 | 6 | 5 | 2 | 3 | -58.34 | 0.058 | 0.062 | 0.026 |
| -1.658 | 0.661 | | | 0.001 | 4 | 6 | 7 | 6 | -59.694 | 0.05 | 0.062 | 0.007 |
| -3.009 | 0.674 | | 0.124 | 0 | 5 | 7 | 6 | 7 | -59.719 | 0.054 | 0.062 | 0.006 |
| 2.811 | | 0.215 | | | 3 | 8 | 9 | 8 | -102.675 | 0.042 | 0.062 | 0 |
| 1.664 | | 0.222 | 0.109 | | 4 | 9 | 8 | 9 | -103.271 | 0.048 | 0.062 | 0 |
| 4.973 | | | | | 2 | 10 | 13 | 10 | -104.809 | 0.038 | 0.062 | 0 |
| 3.994 | | | 0.099 | | 3 | 11 | 12 | 12 | -105.452 | 0.043 | 0.062 | 0 |
| 2.784 | | 0.217 | | 0 | 4 | 12 | 11 | 11 | -105.491 | 0.05 | 0.062 | 0 |
| 1.601 | | 0.23 | 0.105 | 0 | 5 | 13 | 10 | 13 | -106.003 | 0.054 | 0.062 | 0 |
| 4.973 | | | | 0 | 3 | 14 | 15 | 14 | -107.612 | 0.046 | 0.062 | 0 |
| 4.021 | | | 0.095 | 0.001 | 4 | 15 | 14 | 15 | -108.284 | 0.051 | 0.062 | 0 |

Candidate models are ranked by posterior probability ("post. prob."). Ranking in this Bayesian analysis can be compared to ranking via AICc in this analysis and to the ranking in Galipaud et al., 2014 ("rank, post. prob.", *vs.* "rank, AICc" *vs.* "rank (AICc) in Galipaud et al., 2014" columns). First five columns of maximum log-likelihood parameter estimates are part of PyMultinest output. Parameter estimates are reported if present for each of the 16 candidate models. *k,* total number of estimable parameters; *log(Z),* the natural log of the Bayesian evidence/marginal likelihood (*Z*), calculated within the prior-bounded parameter space using our least-squares likelihood function; *log(Z) error,* the error returned by PyMultinest; prior prob., the prior probability that a model is the "correct" model; post. prob., posterior probability that the model is "correct", calculated as $\frac{Z_i * P(i)}{\sum_j Z_j * P(j)}$.

**Table S3. SW and posterior probability calculations for each model variable in both full candidate set and partial candidate set examples.**

| Variable | SW: Galipaud et al. 2014 | SW: this manuscript | SW: candidate subset | Prior probability | Posterior probability | Prior prob: candidate subset | Post prob: candidate subset |
|---|---|---|---|---|---|---|---|
| **x1** | 1 | 1 | 1 | 0.5 | 1 | 0.5 | 1 |
| **x2** | 0.94 | 0.88 | 0.87 | 0.5 | 0.79 | 0.5 | 0.79 |
| **x3** | 0.37 | 0.69 | 0.61 | 0.5 | 0.49 | 0.5 | 0.64 |
| **x4** | 0.37 | 0.25 | 0.09 | 0.5 | 0.07 | 0.5 | 0.07 |

Prior probability for a variable is set at 0.5, meaning a variable's prior probability for can be calculated per candidate model by dividing 0.5 by the number of models in which the variable appears. Prior probability values only impact posterior probability scores and not SW calculations.

### Note S3. SW and posterior probability on a subset of the candidate models.

An important feature of SW is that it is most accurately calculated when the full candidate model set contains equal representation of every variable (1). That is, $x_1$ if appears in 8 models, $x_2$, $x_3$, and $x_4$ must appear in 8 models to calculate an SW for each of them. That is the case in this example from (10). However, we were interested in the case where not every variable may be represented equally in the candidate model set. Once 10 variables are present, a candidate set including equal representation of every variable would include more than 1000 models; more than 13 variables means the full candidate set includes more than 10,000 models. When many variables are present, it is possible to eliminate models from a candidate set, achieving a more tractable number of candidate models, using prior knowledge. We chose to impose synthetic prior knowledge on this 16-model candidate set, as if it is "known" that $x_3$ never appears along with $x_4$. Removing every model that includes both $x_3$ and $x_4$ left us with 12 models rather than 16, where $x_1$ and $x_2$ appeared in 6 models each, while $x_3$ and $x_4$ appeared in 4 models each.

We repeated our analysis on this subset of candidate models (**Table S4**). Similar to our Bayesian analysis of the entire set of candidate models, $x_4$ does not appear until the fifth-highest ranked model; the four highest-ranked models having a cumulative probability of 0.966 (the majority of probability that the "true" model is present); $x_4$ still appears only in candidate models whose posterior probability has decreased compared to the prior (**Table S4**). The SW for $x_4$ in this analysis is 0.09, having changed from its SW of 0.25 using the whole candidate set (**Table S3**). There does not appear to be an accepted threshold over which the SW for a variable indicates it should be included in a model, thus it seems practitioners choose a threshold on a per-case basis (10). As such, it is difficult to assess what a change in SW from 0.25 to 0.09 means when using only a subset of the data – as noted, however, this is not an appropriate use for SWs anyway (1).

**Table S4. Summary of AICc and nested sampling model selection results using a partial candidate set.**

| Model | *log(L)* | AICc | $\Delta_i$ | $w_i$ | Rank, post. prob. | Rank, AICc | *log(Z)* | *log(Z)* error | Prior prob | Post prob |
|---|---|---|---|---|---|---|---|---|---|---|
| $y = x_1 + x_2 + \beta$ | -45.699 | 99.82 | 0 | 0.447 | 0 | 1 | -55.669 | 0.047 | 0.083 | 0.385 |
| $y = x_1 + x_2 + x_3 + \beta$ | -47.526 | 101.301 | 1.482 | 0.213 | 1 | 0 | -55.685 | 0.052 | 0.083 | 0.379 |
| $y = x_1 + \beta$ | -45.724 | 102.085 | 2.266 | 0.144 | 2 | 4 | -56.923 | 0.043 | 0.083 | 0.110 |
| $y = x_1 + x_3 + \beta$ | -47.568 | 103.556 | 3.737 | 0.069 | 3 | 3 | -57.102 | 0.048 | 0.083 | 0.092 |
| $y = x_1 + x_2 + x_4 + \beta$ | -48.778 | 103.807 | 3.987 | 0.061 | 4 | 2 | -58.302 | 0.054 | 0.083 | 0.028 |
| $y = x_1 + x_4 + \beta$ | -50.419 | 104.962 | 5.142 | 0.034 | 5 | 5 | -59.694 | 0.05 | 0.083 | 0.007 |
| $y = x_2 + \beta$ | -48.794 | 106.009 | 6.189 | 0.02 | 6 | 7 | -102.675 | 0.042 | 0.083 | 0 |
| $y = x_2 + x_3 + \beta$ | -50.424 | 107.098 | 7.278 | 0.012 | 7 | 6 | -103.271 | 0.048 | 0.083 | 0 |
| $y = \beta$ | -95.065 | 196.38 | 96.56 | 0 | 8 | 10 | -104.809 | 0.038 | 0.083 | 0 |
| $y = x_3 + \beta$ | -96.334 | 196.791 | 96.971 | 0 | 9 | 9 | -105.452 | 0.043 | 0.083 | 0 |
| $y = x_2 + x_4 + \beta$ | -95.081 | 198.582 | 98.762 | 0 | 10 | 8 | -105.491 | 0.05 | 0.083 | 0 |
| $y = x_4 + \beta$ | -96.336 | 198.921 | 99.102 | 0 | 11 | 11 | -107.612 | 0.046 | 0.083 | 0 |

Candidate models are ranked by posterior probability ("post. prob."). Ranking in this Bayesian analysis can be compared to ranking via AICc in this analysis ("rank, post. prob.", *vs.* "rank, AICc"). Both AICc-related calculations (second through fifth column) and Bayesian calculations (eight through final column) are shown for the partial candidate set results. *log(L)*, maximum log-likelihood; AICc, AIC "corrected" for small sample size; $\Delta_i$, *AICc – min(AICc)* per model; $w_i$, Akaike weight; *log(Z)*, the natural log of the Bayesian evidence/marginal likelihood (*Z*); *log(Z) error*, the error returned by PyMultinest; prior prob., the prior probability that a model is the "correct" model; post. prob., posterior probability that the model is "correct", calculated as $\frac{Z_i * P(i)}{\sum_j Z_j * P(j)}$.

For our Bayesian analysis, the posterior probability that the inclusion of $x_4$ is supported by the data is 0.07, (**Table S3**) and is the same for the full candidate set and for the "prior knowledge excluded" candidate set. Here, $x_4$ is just as unlikely to be included in the "true" model whether assessing a subset or the full set of candidate models. Interestingly, the posterior probability of $x_3$ changes its numerical value when analyzing a subset of the data: from 0.49 in the full candidate set to 0.64 in the “prior knowledge excluded” partial candidate set. However, Bayesian principles dictate that we *can* assess if this is a significant change. With equal prior probability per model variable (**Table S3**), a posterior probability of 0.75 or more, or probability of 0.25 or less, would be considered substantial evidence for inclusion or exclusion of that variable, respectively ((9); see **1.6**). Thus, the change from 0.49 to 0.64 remains in the region between 0.25 and 0.75 where we would consider the data not to have informed whether to include $x_3$ in the model. All other variables ($x_1$, $x_2$, $x_4$) remained at their same values and thus remained either in the “informed and should be included” set ($x_1$, $x_2$) or the “informed and should be excluded” set ($x_4$). It is likely that the weak correlation of $x_3$ with the response variable resulted in the Bayesian analysis result of “unsure” whether $x_3$ should be included.

**Table S5. Existing data pertaining to SCLC intratumoral heterogeneity and communication used for rate parameter priors.**

| |
|---|
| NCI-H69 cell line (SCLC-A) doubles at 51.1 +/- 3.1 hours, equivalent to 0.469 doublings per day (13). |
| NCI-H82 (SCLC-N) doubles at 25.5 +/- 4.2 hours, equivalent to 0.783 doublings per day (13). |
| NCI-H841 (SCLC-Y) doubles at 31.2 +/- 3.6 hours, equivalent to 0.769 doublings per day (13). |
| DMS53 (SCLC-A2) doubles at ~127 hours, equivalent to 0.1898 doublings per day (14). |
| Average of apoptotic indices for SCLC cell lines is 0.081, used as 0.081 deaths per day (15). |
| Little is known about timing of phenotypic transitions so a very permissive range was used in the model, with values considered uniformly likely from 0.01 transitions per day (1 transition every ~3 months) to 3 transitions per day. We find this to be a reasonable permissive range based on mechanistic modeling of the epithelial-to-mesenchymal transition in breast cancer cells and stem cell differentiation (16,17), as transition rates for SCLC subtypes have not been reported in the literature. In these reports, the EMT transition was fit to between 10 (16) and 20 (17) days. |
| Changes in growth rates specifically due to inter-subtype effects have not been recorded. NE viability (luciferase) and division (EdU incorporation) are increased by NonNE when plated together (18); NonNE growth decreases in the presence of NE (19). We used a 5% increase or decrease of the baseline value (depending on subtype and interaction) as the parameter prior for each affected rate. |

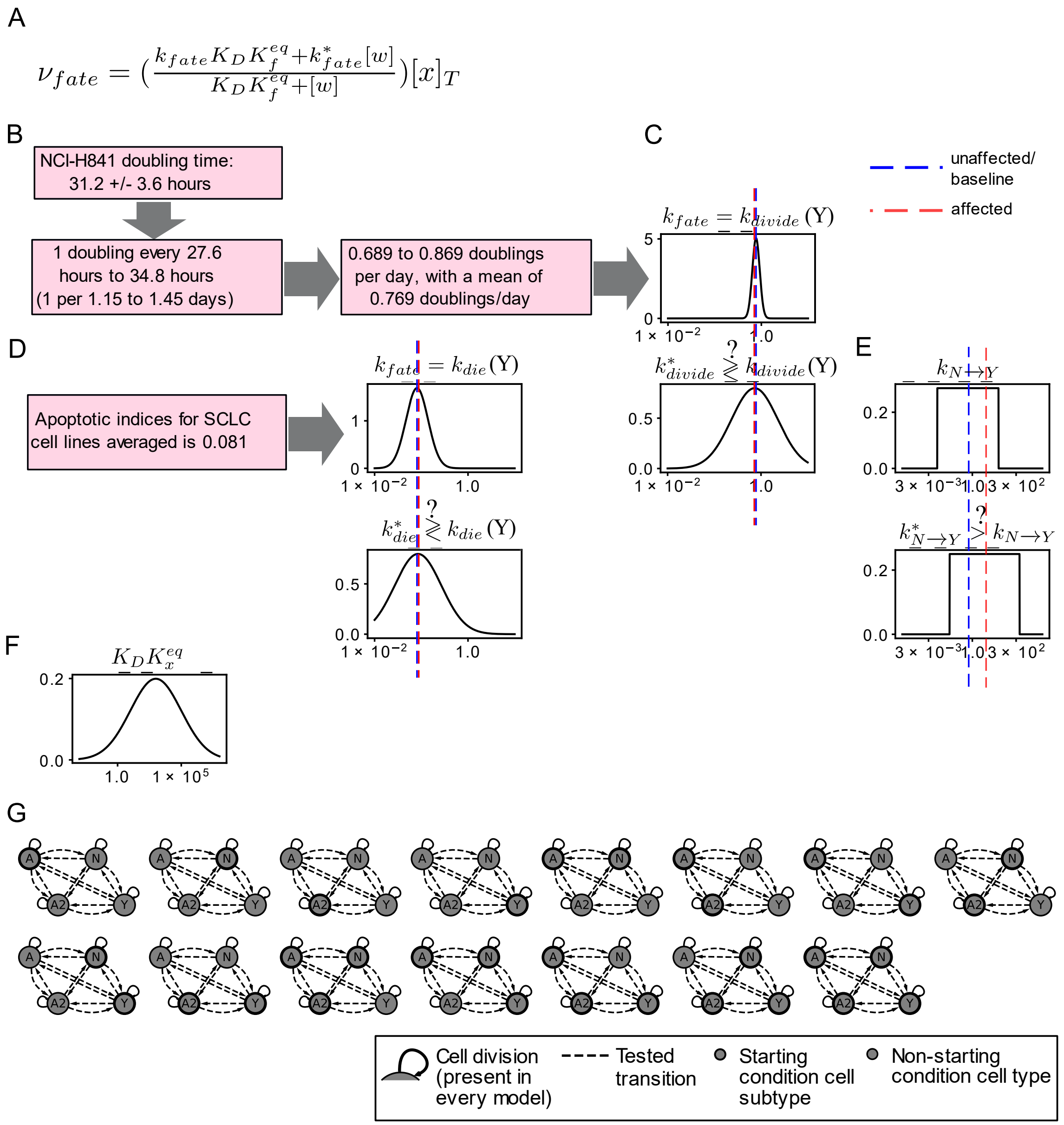


**Fig S2. Prior probabilities values and schematics.** (A) Rate of a cell fate (division, death, or phenotypic transition) for *x* ($v_{fate}$) can be calculated as a function of the population size of the effector cell *w* (see **Box 1**) [21]. (B) Example calculation of division rate parameter prior for H841, representation of subtype Y, (see **Table S5**) converting doubling times to "per day" units. (C) Division prior for subtype Y, (blue dashed line centered at the mean) as well as inter-subtype effect on division, whose mean is centered 5% lower (red dashed line; see **Table S5**) with wider variance to account for more uncertainty in inter-subtype effects. (D) Example calculation and visualization of death rate parameter prior for Y (blue dashed line at mean) and inter-subtype effect on death (red dashed line at mean, 5% higher). (E) Example uniform transition prior, (see **Table S5**) here showing N to Y transition; blue dashed line at baseline transition rate center, red dashed line at inter-subtype effect transition rate center. (F) Equilibrium assumption prior, representing $K_D K_x^{eq}$ in the equation (A). Each affected interaction has a unique $K_D K_x^{eq}$ prior, but all such priors have identical values (centered at 1000) before fitting. (G) Different model initiation hypotheses, where a model can be initiated by one or more subtypes (thick black outline) depending on the subtypes present (four-subtype topology shown). With 15 options and equal prior probabilities, the prior probability for each initiation hypothesis is 6.67%.

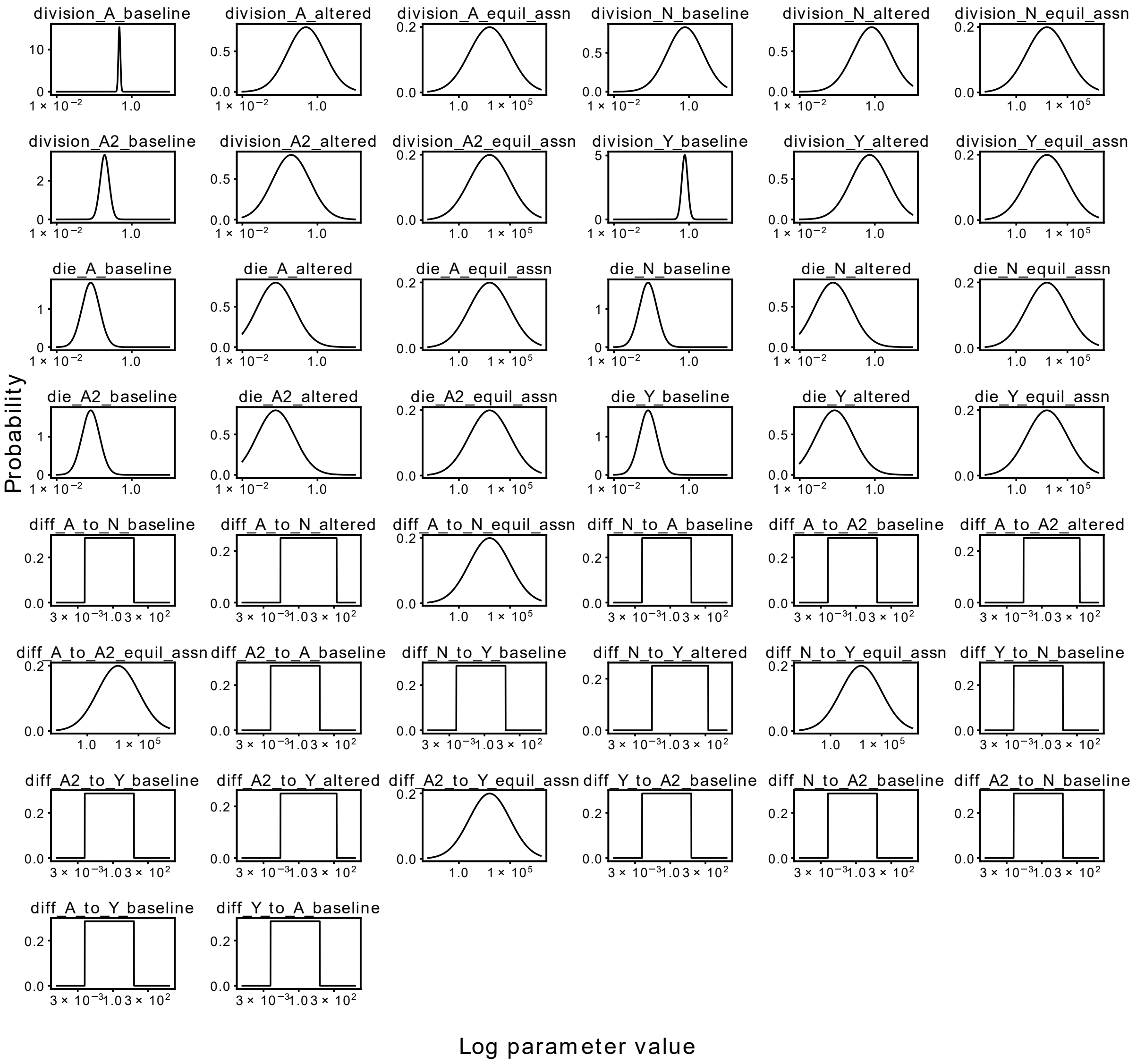


**Fig S3. Parameter prior distributions for all possible reactions in a candidate population dynamics model.** If a candidate model does not contain a reaction, for example a model with the topology A, N, and Y does not include A2 and thus will not include A2 division, death, or transitions to/from A2, then the rate parameter priors for A2-related reactions will not be included as a parameter prior for model fitting.

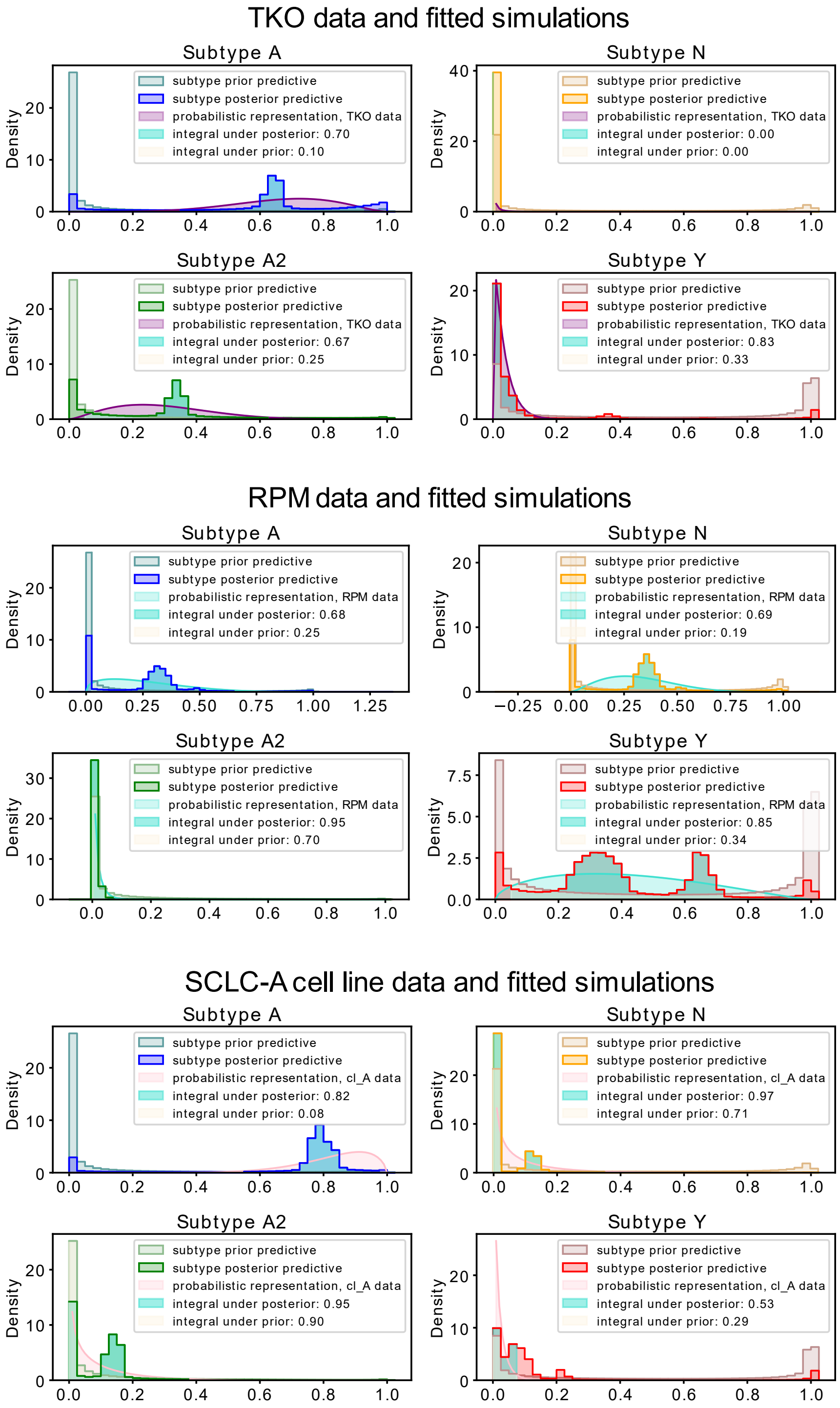


**Fig S4. Nested sampling's fitting results in better-fitting simulations than simulations using randomly selected parameter values.** Data distribution prior predictive distribution, and posterior predictive distribution for each dataset and all candidate models. Data is fit to a Beta distribution, bounded by zero and one, and used in the likelihood function input for Multinest (see **Methods**). Prior predictive distribution represents model simulations using parameters randomly drawn from the prior. Posterior predictive is generated by model simulations using best-fitting parameters returned by Multinest. See **Note S5** for more detail.

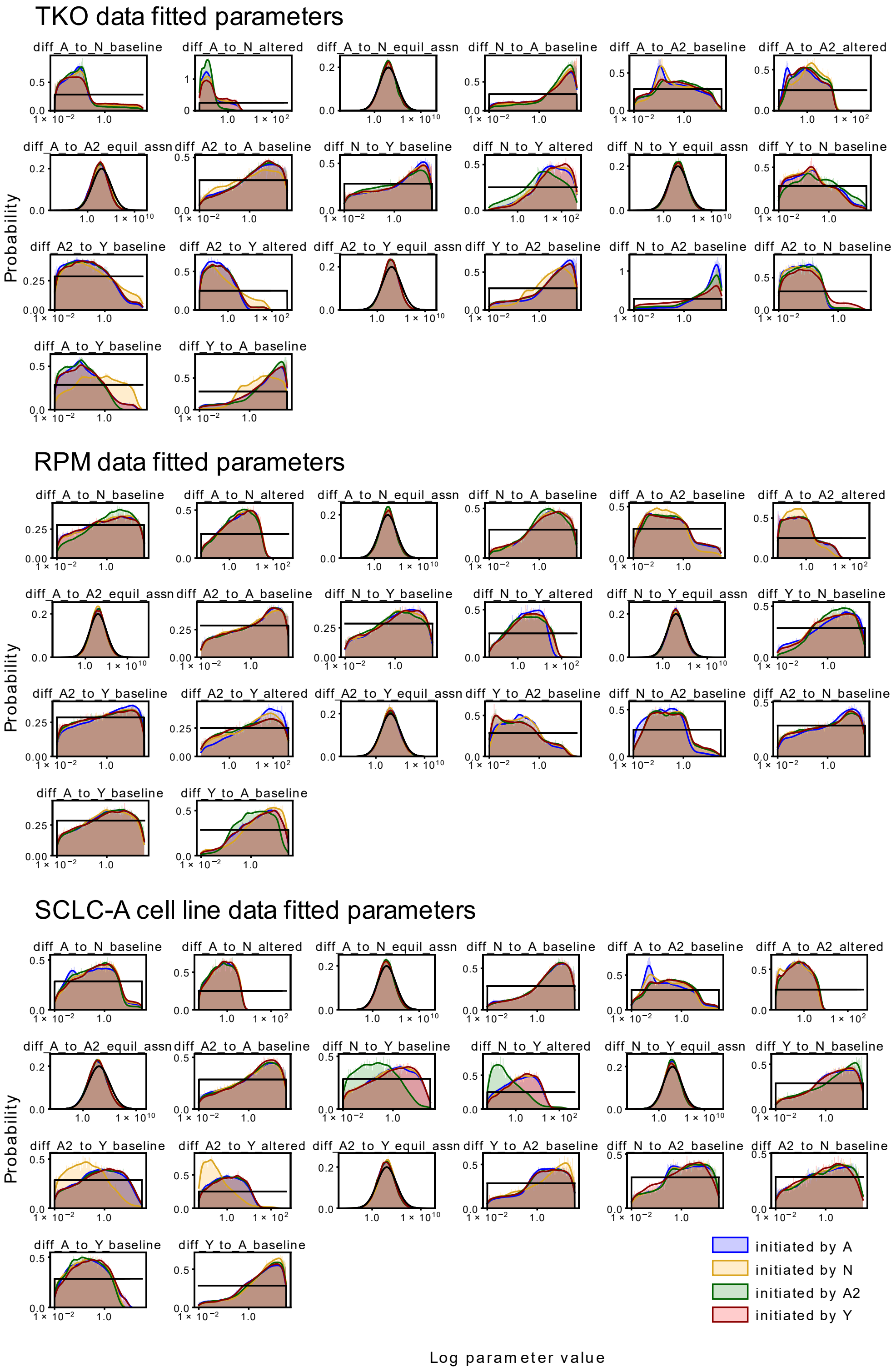


**Fig S5. Rate parameter posterior marginal distributions, after applying modeling averaging based on candidate model posterior probability.** Rate parameter distributions are plotted based on candidate model initiating subtype, where the subtype that initiates the tumor in the simulation results in nonidentical posterior distributions for some model parameters.

**Note S4. All datasets support alteration of phenotypic transition rates in the presence of N or A2 subtypes**

The highest likelihood model topologies (**Fig 4A**, blue) for the TKO GEMM data, along with the four-subtype topology, are compared in **Fig 4B** (left). Three model variables have significantly different parameter rates across model topologies: (*i*) the A-to-Y transition, (*ii*) the Y-to-A transition; and *(iii*) the A-to-A2 transition. The A-to-Y transition has a slower rate if A2 is present in the population and the Y-to-A transition has a faster rate if A2 is present. However, the presence of N along with A2 does not change the rate of either transition. Similarly, only Y affects the A-to-A2 transition, increasing its rate. N does decrease the rate of the A2-to-A transition despite no effect from Y. The Y-to-A2 transition rate is an increased in the presence of A2. These observations have mechanistic implications: A2 may represent an intermediate subpopulation in the tumor that is longer-lived, and will only slowly transition to Y. In the topology with A, A2, and Y (**Fig 4A**, structure 2), the A-to-A2 transition takes up more of the flux in the network. Additionally, the N-to-Y transition is faster relative to the A2-to-Y transition (**Fig S6B**), suggesting that N is a shorter-lived intermediate in the A-to-N-to-Y transition. This result aligns with previous experiments (20) where N was identified as a short-lived state in the A-to-N-to-Y transition. We therefore predict that A2 and N are involved in regulating the relative abundance of, and flux between, A and Y in the tumor.

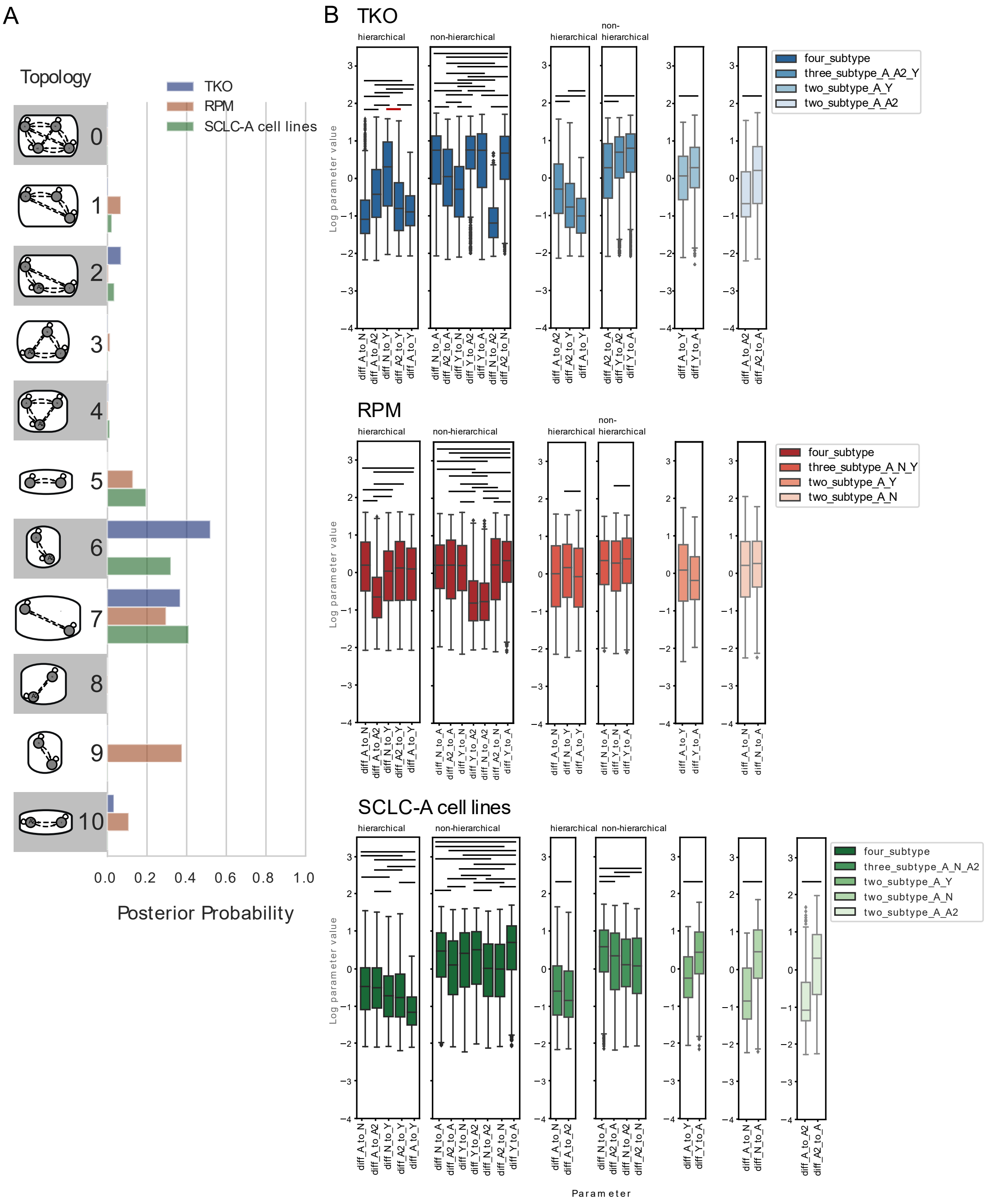


**Fig S6. Transition parameter rates vary in similar ways across datasets.** (A) Hypothesis assessment of model topologies per dataset, posterior probabilities based on all candidate models, with no filtering based on initiating subtype (see **Fig S4**). Model topologies represented by images and corresponding numbers along the y-axis. (B) Comparison of phenotypic transition parameter posterior marginal distributions, BMA-weighted, per dataset, separated by topology. In 3- and 4-subtype topologies, distributions are further separated by hierarchical or non-hierarchical transition status. Bars indicate significance between samples from BMA parameter distributions at family-wise error rate (FWER) of 0.01, using one-way ANOVA plus Tukey HSD. Red bar: comparison noted in the main text.

We also compared the highest likelihood model topologies (**Fig 4A**, red) for RPM-fitted models, as well as the four-subtype topology (**Fig 4B**, middle). Five parameter rates are significantly different across model topologies. These are (*i*) the A-to-Y transition, (*ii*) the Y-to-A transition; (*iii*) the A-to-N transition; *(iv)* the N-to-Y transition; and *(v)* the N division rate. The same transitions affected in the TKO-fitted models (A-to-Y and Y-to-N transitions) are affected in the RPM models in the same ways, despite the experimental data being different. We thus predict that N and A2 are modulating the transition between, and relative abundance of, A and Y. Unlike in the TKO data, when A2 is present the flux through the system spends more time in the N subtype, with more frequent transitions to N and less frequent transitions to Y; additionally, the rate of the A2-to-N transition is faster than the N-to-A2 transition (**Fig S6B**). We predict that while N may be a shorter-lived intermediate than A2, A2 regulates the flux from A-to-N-to-Y.

Next, we compared the highest likelihood model topologies (**Fig 4A**, green) for the SCLC-A cell line data and the four-subtype topology (**Fig 4B**, middle). Six parameter rates are significantly different across model topologies, five of which recapitulate rate alterations based on the presence or absence of different subtypes in TKO or RPM datasets. The rate alteration unique to the SCLC-A dataset is the A2-to-A transition, which is less frequent in the four-subtype topology, indicating that the presence of Y decreases this rate. However, in the TKO dataset, the A2-to-A transition decreases with addition of N.

**Note S5. Simulations using best-fitted parameters, as opposed to randomly-selected parameters from the prior distributions, replicate subtype proportions at steady state**

Each different dataset (different model selection run) differentiated between more likely and less likely models for that particular dataset. Given that model selection determines Bayesian evidence scores by balancing model complexity with goodness of fit, we expected that the subtype proportions at steady state in the highest scoring models would replicate cell subtype proportions in the data.

Simulating all fitted models, using parameters selected only from the prior marginal distributions, the subtype proportions at steady state for each subtype tended to fall at 0 (0% of the simulated tumor) or 1 (100% of the simulated tumor), indicating that parameters chosen at random from the prior distributions do not fit the data (**Fig S4**). However, in selecting parameters from the posterior, fitted, distributions – those representing the highest-scoring parameter sets – the simulations matched the data much more closely. There still remained simulations where the subtypes fell either at 0% or 100% of the simulated tumor, but the subtype steady state fell within the probabilistic representation of the data more of the time (**Fig S4**).

For the TKO dataset, not taking into account subtype N, for which all simulations both using prior parameters and fitted parameters resulted in a steady state of 0%, on average 23% of the prior predictive density fell within the 95% confidence interval of the probabilistic representation of the data (**Fig S4**, purple). After parameter fitting and evidence calculation, on average 73% of the posterior predictive density fell within the 95% confidence interval of the data. (Since the data, the prior predictive, and the posterior predictive are all probabilistic densities, each curve integrates to 1 or 100%.) Investigating these numbers it is clear that the model selection / parameter fitting process was unable to bring simulated subtype steady states completely within the 95% confidence interval of the TKO data subtype proportions, though it has brought it closer than the prior predictive. We would interpret this as the fact that the TKO data used for fitting was not able to completely outweigh the subtype proportions resulting from simulations using the prior parameters, which is a known possibility with regard to using Bayes' Theorem to fit a model's parameter sets to data (12).

For the RPM dataset, not taking into account subtype A2, for which all simulations both using prior parameters and fitted parameters resulted in a steady state of 0%, on average 26% of the prior predictive density fell within the 95% confidence interval of the probabilistic representation of the data. After parameter fitting and evidence calculation, on average 74% of the posterior predictive density fell within the 95% confidence interval of the data. Interestingly, the posterior predictive density for the Y subtype was bimodal (**Fig S4**, middle), which we expect is related to the very wide range of the probabilistic representation of Y proportions in this dataset (**Fig 2A**). For the SCLC-A clustered cell lines dataset, on average 50% of the prior predictive density fell within the 95% confidence interval of the probabilistic representation of the data. After parameter fitting and evidence calculation, on average 82% of the posterior predictive density fell within the 95% confidence interval of the data. In all, simulations using fitted parameters regardless of dataset resulted in a 30-50% better correspondence to subtype proportions in the data, indicating that the process of model selection and model averaging resulted in models and parameter sets that were able to represent the data at hand to a satisfactory extent.

| Table S6. Model term posterior probabilities after hypothesis exploration, TKO high-probability 3-subtype topology | | | |
|---|---|---|---|
| **Model variable** | **Candidate model prior per hypothesis / summed prior** | **Model-averaged posterior probability** | **Odds ratio** |
| A to N transition | N/A | N/A | N/A |
| A to A2 transition | $P(M\|H_{A\to A2}) = 0.0008$, $P(M\|H_{no}) = 0.006$ / sum 0.5 vs 0.5 | $P(H_{A\to A2}\|D) = 0.62$ | 1.63 |
| N to Y transition | N/A | N/A | N/A |
| A2 to Y transition | $P(M\|H_{A2\to Y}) = 0.0008$, $P(M\|H_{no}) = 0.004$ | $P(H_{A2\to Y}\|D) = 0.51$ | 1.04 |
| A to Y transition | $P(M\|H_{A\to Y}) = 0.001$, $P(M\|H_{no}) = 0.003$ / sum 0.5 vs 0.5 | $P(H_{A\to Y}\|D) = 0.64$ | 1.78 |
| N to A2 transition | N/A | N/A | N/A |
| A2 to N transition | N/A | N/A | N/A |
| N to A transition | N/A | N/A | N/A |
| A2 to A transition | $P(M\|H_{A2\to A}) = 0.002$, $P(M\|H_{no}) = 0.001$ / sum 0.5 vs 0.5 | $P(H_{A2\to A}\|D) = 0.71$ | 2.45 |
| Y to N transition | N/A | N/A | N/A |
| Y to A2 transition | $P(M\|H_{Y\to A2}) = 0.0015$, $P(M\|H_{no}) = 0.0013$ / sum 0.5 vs 0.5 | $P(H_{Y\to A2}\|D) = 0.68$ | 2.13 |
| Y to A transition | $P(M\|H_{Y\to A}) = 0.0018$, $P(M\|H_{no}) = 0.0011$ / sum 0.5 vs 0.5 | $P(H_{Y\to A}\|D) = 0.81$ | 4.26 |
| Non-NE affects division & death | $P(M\|H_{div\_eff}) = 0.0012$, $P(M\|H_{no}) = 0.0016$ / sum 0.5 vs 0.5 | $P(H_{div\_eff}\|D) = 0.44$ | 0.78 |
| Y affects division & death vs A2&Y affect division & death | $P(M\|H_{div\_eff\_Y}) = 0.0016$, $P(M\|H_{div\_eff\_A2\_Y}) = 0.0016$, $P(M\|H_{no}) = 0.0011$ / sum 0.33 vs 0.33 vs 0.33 | $P(H_{div\_eff\_Y}\|D) = 0.28$, $P(H_{div\_eff\_A2\_Y}\|D) = 0.33$ | 0.39, 0.50 |
| Non-NE affects early transitions (A-N, A-A2) | $P(M\|H_{early\_eff}) = 0.0011$, $P(M\|H_{no}) = 0.0018$ / sum 0.5 vs 0.5 | $P(H_{early\_eff}\|D) = 0.35$ | 0.54 |
| Y affects early transitions (A to N, A to A2) vs A2&Y affect these | $P(M\|H_{early\_eff\_Y}) = 0.0015$, $P(M\|H_{early\_eff\_A2\_Y}) = 0.0015$, $P(M\|H_{no}) = 0.0012$ / sum 0.33 vs 0.33 vs 0.33 | $P(H_{early\_eff\_Y}\|D) = 0.24$, $P(H_{early\_eff\_A2\_Y}\|D) = 0.28$ | 0.32, 0.39 |
| Non-NE affects late transitions (N-Y, A2-Y) | $P(M\|H_{late\_eff}) = 0.0021$, $P(M\|H_{no}) = 0.0010$ / sum 0.5 vs 0.5 | $P(H_{late\_eff}\|D) = 0.29$ | 0.41 |
| Y affects late transitions (N to Y, A2 to Y) vs A2&Y affect these | $P(M\|H_{late\_eff\_Y}) = 0.0028$, $P(M\|H_{late\_eff\_A2\_Y}) = 0.0027$, $P(M\|H_{no}) = 0.0007$ / sum 0.33 vs 0.33 vs 0.33 | $P(H_{eff\_from\_Y}\|D) = 0.21$, $P(H_{eff\_from\_A2}\|D) = 0.24$ | 0.27, 0.32 |
| If Non-NE effect true, comes from Y or A2&Y? | $P(M\|H_{eff\_from\_Y}) = 0.0015$, $P(M\|H_{eff\_from\_A2\_Y}) = 0.0015$ / sum 0.5 vs 0.5 | $P(H_{eff\_from\_Y}\|D) = 0.47$, $P(H_{eff\_from\_A2\_Y}\|D) = 0.53$ | 0.89, 1.13 |

| Table S7. Model term posterior probabilities after hypothesis exploration, RPM high-probability 3-subtype topology | | | |
|---|---|---|---|
| **Model variable** | **Candidate model prior per hypothesis / summed prior** | **Model-averaged posterior probability** | **Odds ratio** |
| A to N transition | $P(M\|H_{A\to N}) = 0.0014$, $P(M\|H_{no}) = 0.0086$ / sum 0.5 vs 0.5 | $P(H_{A\to N}\|D) = 0.71$ | 2.45 |
| A to A2 transition | N/A | N/A | N/A |
| N to Y transition | $P(M\|H_{N\to Y}) = 0.0015$, $P(M\|H_{no}) = 0.0068$ / sum 0.5 vs 0.5 | $P(H_{N\to Y}\|D) = 0.6$ | 1.50 |
| A2 to Y transition | N/A | N/A | N/A |
| A to Y transition | $P(M\|H_{A\to Y}) = 0.0017$, $P(M\|H_{no}) = 0.0042$ / sum 0.5 vs 0.5 | $P(H_{A\to Y}\|D) = 0.71$ | 2.45 |
| N to A2 transition | N/A | N/A | N/A |
| A2 to N transition | N/A | N/A | N/A |
| N to A transition | $P(M\|H_{N\to A}) = 0.0027$, $P(M\|H_{no}) = 0.0022$ / sum 0.5 vs 0.5 | $P(H_{N\to A}\|D) = 0.78$ | 3.55 |
| A2 to A transition | N/A | N/A | N/A |
| Y to N transition | $P(M\|H_{Y\to N}) = 0.0028$, $P(M\|H_{no}) = 0.0022$ / sum 0.5 vs 0.5 | $P(H_{Y\to N}\|D) = 0.68$ | 2.13 |
| Y to A2 transition | N/A | N/A | N/A |
| Y to A transition | $P(M\|H_{Y\to A}) = 0.0033$, $P(M\|H_{no}) = 0.0020$ / sum 0.5 vs 0.5 | $P(H_{Y\to A}\|D) = 0.81$ | 4.26 |
| Non-NE affects division & death | $P(M\|H_{div\_eff}) = 0.0025$, $P(M\|H_{no}) = 0.0024$ / sum 0.5 vs 0.5 | $P(H_{div\_eff}\|D) = 0.46$ | 0.85 |
| Y affects division & death vs A2&Y affect division & death | N/A (since A2 as part effect not possible, calculations are same as Non-NE affects division, death, above) | N/A | N/A |
| Non-NE affects early transitions (A-N, A-A2) | $P(M\|H_{early\_eff}) = 0.0023$, $P(M\|H_{no}) =$ 0.0027 / sum 0.5 vs 0.5 | $P(H_{early\_eff}\|D) = 0.44$ | 0.79 |
| Y affects early transitions (A to N, A to A2) vs A2&Y affect these | N/A (since A2 as part effect not possible, calculations are same as Non-NE affects early transitions, above) | N/A | N/A |
| Non-NE affects late transitions (N-Y, A2-Y) | $P(M\|H_{late\_eff}) = 0.0042$, $P(M\|H_{no}) = 0.0017$ / sum 0.5 vs 0.5 | $P(H_{late\_eff}\|D) = 0.37$ | 0.59 |
| Y affects late transitions (N to Y, A2 to Y) vs A2&Y affect these | N/A (since A2 as part effect not possible, calculations are same as Non-NE affects late transitions, above) | N/A | N/A |
| If Non-NE effect true, comes from Y or A2&Y? | N/A (A2 as part of the Non-NE effect not possible) | N/A | N/A |

| **Table S8. Model term posterior probabilities after hypothesis exploration, SCLC-A cell line data high-prob. 3-subtype topology** | | | |
|---|---|---|---|
| **Model variable** | **Candidate model prior per hypothesis / summed prior** | **Model-averaged posterior probability** | **Odds ratio** |
| A to N transition | $P(M\|H_{A\to N}) = 0.0031$, $P(M\|H_{no}) = 0.011$ / sum 0.5 vs 0.5 | $P(H_{A\to N}\|D) = 0.61$ | 1.56 |
| A to A2 transition | $P(M\|H_{A\to A2}) = 0.0031$, $P(M\|H_{no}) = 0.011$ / sum 0.5 vs 0.5 | $P(H_{A\to A2}\|D) = 0.52$ | 1.10 |
| N to Y transition | N/A | N/A | N/A |
| A2 to Y transition | N/A | N/A | N/A |
| A to Y transition | N/A | N/A | N/A |
| N to A2 transition | $P(M\|H_{N\to A2}) = 0.0038$, $P(M\|H_{no}) = 0.0064$ / sum 0.5 vs 0.5 | $P(H_{N\to A2}\|D) = 0.66$ | 1.94 |
| A2 to N transition | $P(M\|H_{A2\to N}) = 0.0038$, $P(M\|H_{no}) = 0.0064$ / sum 0.5 vs 0.5 | $P(H_{A2\to N}\|D) = 0.66$ | 1.94 |
| N to A transition | $P(M\|H_{N\to A}) = 0.0058$, $P(M\|H_{no}) = 0.0041$ / sum 0.5 vs 0.5 | $P(H_{N\to A}\|D) = 0.74$ | 2.85 |
| A2 to A transition | $P(M\|H_{A2\to A}) = 0.0058$, $P(M\|H_{no}) = 0.0041$ / sum 0.5 vs 0.5 | $P(H_{A2\to A}\|D) = 0.61$ | 1.56 |
| Y to N transition | N/A | N/A | N/A |
| Y to A2 transition | N/A | N/A | N/A |
| Y to A transition | N/A | N/A | N/A |
| Non-NE affects division & death | $P(M\|H_{div\_eff}) = 0.005$, $P(M\|H_{no}) = 0.0046$ / sum 0.5 vs 0.5 | $P(H_{div\_eff}\|D) = 0.16$ | 0.19 |
| Y affects division & death vs A2&Y affect division & death | N/A (since Y effect not possible, calculations are the same as Non-NE affects division & death, above) | N/A | N/A |
| Non-NE affects early transitions (A-N, A-A2) | $P(M\|H_{early\_eff}) = 0.0051$, $P(M\|H_{no}) = 0.0045$ / sum 0.5 vs 0.5 | $P(H_{early\_eff}\|D) = 0.17$ | 0.21 |
| Y affects early transitions (A to N, A to A2) vs A2&Y affect these | N/A (since Y effect not possible, calculations are the same as Non-NE affects early transitions, above) | N/A | N/A |
| Non-NE affects late transitions (N-Y, A2-Y) | N/A (Y not in this model so no transitions toward it) | N/A | N/A |
| Y affects late transitions (N to Y, A2 to Y) vs A2&Y affect these | N/A (Y not in this model so no transitions toward it) | N/A | N/A |
| If Non-NE effect is true, comes from Y or A2&Y? | N/A (Y effect not possible) | N/A | N/A |
